# Methylation pseudotime analysis for label-free profiling of the temporal chromatin landscape with long-read sequencing

**DOI:** 10.1101/2025.03.03.641287

**Authors:** Annie Trinh, Navied Akhtar, Kwadwo Bonsu, Nandor Laszik, Asia Mendelevich, Tanye Wen, Julien L. P. Morival, Katelyn E. Diune, Mitchell Frazeur, Justin E. Vega, Alexander A. Gimelbrant, Elizabeth L. Read, Timothy L. Downing

## Abstract

Faithful epigenetic inheritance across cell divisions is essential to maintaining cell identity and involves numerous epigenetic modifications, whose roles in establishing chromatin architecture are less understood. Technological approaches to temporally order epigenetic modifications throughout the cell cycle often face limitations in sequence resolution and rely on potentially damaging mitotic labeling or conversion steps. Herein, we present <u>M</u>ethylation <u>P</u>seudotime <u>A</u>nalysis <u>T</u>hrough read-level <u>H</u>eterogeneity (MPATH), a label- and conversion-free method to infer post-replication DNA strand maturity from methylation patterns across single molecules. We use MPATH to temporally order hydroxymethylation throughout mitotic inheritance, revealing that CpGs within cis-regulatory elements undergo transitions between methylation states at sub-cell-cycle timescales. When applied to long reads generated by NOMe-seq, MPATH uncovered relationships between nucleosome occupancy and DNA maturity. Finally, extension of MPATH to phased reads reveals allele-specific trends in pseudotime distribution associated with X chromosome activity. Our findings suggest that when coupled with multimodal sequencing strategies, MPATH could provide valuable insights into chromatin restoration dynamics.

## Main

While individual cells of a multicellular organism carry the same genetic code, the establishment and maintenance of unique cell identities is made possible by epigenetic regulation of gene expression^1,2^. To maintain cell identity following genome replication, the epigenetic landscape must be faithfully restored to nascent DNA^3^. Chromatin re-establishment is an orchestrated process involving numerous epigenetic modifications and modifiers, whose functional roles in collaboratively shaping chromatin architecture are not well understood^4,5^. Given that the epigenome needs to be de- and reassembled during each cycle of replication, the ordering of DNA modifications across sub-cell-cycle timescales could inform the collaborative processes underlying epigenetic crosstalk.

While time-lapse microscopy and mass spectrometry have enabled tracking of epigenetic modifications throughout time^6–9^, these approaches often lack base-pair resolution, which may be valuable for understanding sequence-specific contexts of epigenetic inheritance. To achieve temporal resolution of epigenetic modifications through sequencing, we previously applied mitotic labeling in human embryonic stem cells (hESCs) to track CpG methylation on nascent DNA and reported a global delay in CpG remethylation following genome replication, which can be attributed to kinetic heterogeneity across CpG sites^10,11^. Other studies that employed mitotic labeling and bisulfite sequencing reported that replication-uncoupled CpG methylation can be impeded by nucleosome occupancy, suggesting that the replication-associated dynamics of epigenetic modifiers and modifications could inform how epigenetic marks propagate and influence each other throughout time^12^. While current replication-coupled sequencing approaches typically capture epigenomic data at length scales typically under 250 base pairs (bp), long-read sequencing has uncovered correlations between CpG methylation and nucleosome positioning that are otherwise masked at bulk resolution^13^. Also importantly, mitotic labeling with nucleoside analogs can be cytotoxic and influence cell identity^14–16^, while bisulfite conversion can induce unwanted DNA degradation^17–19^.

Herein, we present <u>M</u>ethylation <u>P</u>seudotime <u>A</u>nalysis <u>T</u>hrough read-level <u>H</u>eterogeneity (MPATH), a label- and conversion-free method for determining DNA strand maturity with respect to genome replication. When coupled with other multimodal sequencing strategies, MPATH enables the temporal ordering of epigenetic modifications across sub-cell-cycle timescales with long-read and single-molecule resolution. We first demonstrated with BrdU labeling that nascent and mature chromatin can be distinguished by intramolecular CpG methylation patterns across individual long reads generated by nanopore sequencing. Then, we leveraged these patterns to infer chromatin maturity in unlabeled long reads. Benchmarking results demonstrated that MPATH can recapitulate previously observed CpG remethylation dynamics without DNA labeling, eliminating the need for nucleoside analogs and bisulfite conversion. We used MPATH to temporally order CpG hydroxymethylation across replication-associated timescales in hESCs and identified a subset of CpGs that exhibit a putative switch-like behavior between hydroxymethylated and unhydroxymethylated states at cis-regulatory elements. When applied to existing long-read Nucleosome Occupancy and Methylome-sequencing (NOMe-seq) datasets, MPATH revealed an enrichment of mononucleosomes in nascent chromatin which was lost over time. We also applied MPATH to phased long reads from Abl.1 mouse B cells and observed that X chromosome alleles can progress differently across replication-associated timescales according to activity. Our findings suggest that when coupled with ectopic and intrinsic DNA modification mapping strategies, MPATH can dissect the molecular underpinnings of dynamic, multi-factor chromatin restoration.

## Results

### Nanopore long-read sequencing enables detection and tracking of nascent strand DNA methylation across sub-cell-cycle timescales

Following genome replication, the dynamics of epigenetic modifications underlying chromatin maturation are yet to be fully elucidated (Fig. 1a). Given that global CpG methylation is restored with a pronounced and highly variable delay in hESCs (Supplemental Fig. 1a), we sought to leverage intramolecular CpG methylation patterns toward developing a label-free method that would enable the temporal ordering of individual long reads (Fig. 1b). We posited that such a method, when coupled with DNA modification mapping strategies, would be effective toward profiling the dynamics of epigenetic modifications and modifiers across replication-associated timescales. To demonstrate the utility of nanopore long-read sequencing in detecting cell cycle-associated methylation states, we induced cell cycle arrest in HUES64 hESCs using 2 µg/mL nocodazole. Comparison of read-level methylation fraction (fraction of CpGs that are methylated across single reads) to bulk methylation fraction (average bulk methylation of the individual CpGs corresponding to those in a given read, derived from whole genome bisulfite sequencing, or WGBS) revealed an increasing shift in read-level methylation states upon cell cycle arrest, reflecting the suitability of the nanopore long-read sequencing platform for detecting cell cycle-associated methylation states (Fig. 1c).

**Figure 1.**
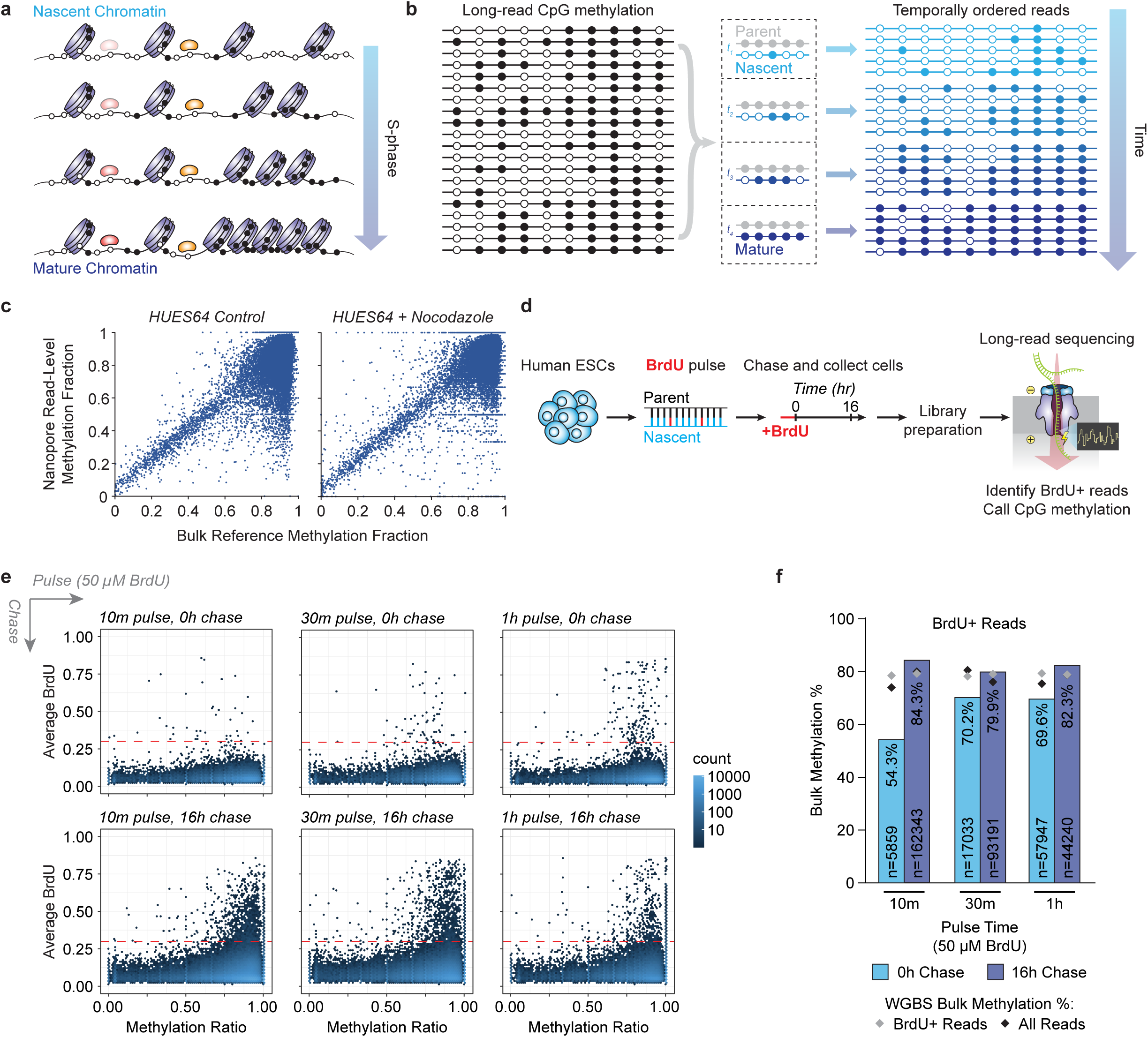
ONT long-read sequencing enables detection and tracking of nascent strand DNA methylation across sub-cell-cycle timescales. **a)** Illustrative schematic of chromatin maturation across S-phase. CpG sites are represented as circles (black: methylated, white: unmethylated). Different temporal dynamics are shown for transcription factors in red and orange. Purple nucleosomes are shown to exhibit shifts in positioning throughout time. **b)** Illustrative schematic of methylation pseudotime analysis. Individual long reads can be temporally ordered according to their intramolecular CpG methylation patterns. **c)** Scatter plots comparing read-level methylation fraction for individual long reads to corresponding bulk reference methylation fraction derived from WGBS between control and nocodazole-treated HUES64 hESCs. **d)** Schematic workflow of BrdU pulse-chase experiment, followed by ONT long-read sequencing and detection of modified bases. **e)** Hexbin plots depicting methylation ratio and average BrdU content across individual long reads generated from pulse optimization experiments. Red dashed line corresponds to an average BrdU threshold derived from a mock-treated negative control. Reads with average BrdU content above this threshold are considered BrdU-positive. **f)** Bar graph depicting bulk methylation percentage of BrdU-positive reads from BrdU-labeled experimental conditions, as calculated by total methylated CpGs divided by total CpGs. Gray diamond represents the bulk methylation percentage for reads generated by WGBS that capture the corresponding CpGs in BrdU-positive long reads from a given condition. Black diamond represents the bulk methylation percentage for reads generated by WGBS that capture the corresponding CpGs in all long reads from a given condition.

To characterize how nascent and mature chromatin differ in their intramolecular CpG methylation states, we applied mitotic labeling in hESCs and performed nanopore sequencing (Fig. 1d). Briefly, HUES64 hESCs were pulsed with 50 µM BrdU to label replicating DNA and immediately collected (0h timepoint) and after 16h chase (corresponding to nascent and mature chromatin, respectively) before sequencing. BrdU-containing reads and CpG methylation were detected using nucleotide analog- and methylation-sensitive basecalling, respectively (Fig. 1e, Supplemental Fig. 1b,c, and Supplemental Table 1) (Methods). On average, sequenced reads generated 1.3x coverage per sample, with an average read length of 5.8 kb (Supplemental Table 1). We observed that CpGs from BrdU-containing reads collected at the 0h chase timepoint were hypomethylated relative to the 16h chase timepoint and bulk CpG methylation levels obtained from WGBS, supporting previous observations of a global delay in nascent DNA methylation (Fig. 1f). While increasing the pulse duration enabled more BrdU integration and therefore more BrdU-containing reads, it also resulted in a tradeoff in our ability to capture nascent (versus mature) DNA methylation levels (Fig. 1f).

To obtain a higher fraction of BrdU-labeled reads while still capturing nascent methylation, we performed a second pulse-chase experiment using a 10-minute pulse time with a 10-fold increase in BrdU concentration (500 µM) (Supplemental Table 1). Notably, cell viability remained largely unchanged across a range of BrdU concentrations up to 100-fold across this short pulse duration (Supplemental Fig. 1d). Additionally, BrdU integration did not appear to strongly favor particular genomic regions (Supplemental Fig. 1e). Across both replicates that were pulsed with 500 µM BrdU for 10 minutes, we generated an average of 2x coverage per sample, with an average N50 read length of 7.6 kb (Supplemental Table 1). Replicates captured a similar delay in nascent DNA methylation levels (23.6% and 24.5% difference in bulk methylation of BrdU-containing reads between 0 and 16h chase timepoints) (Fig. 2a).

**Figure 2.**
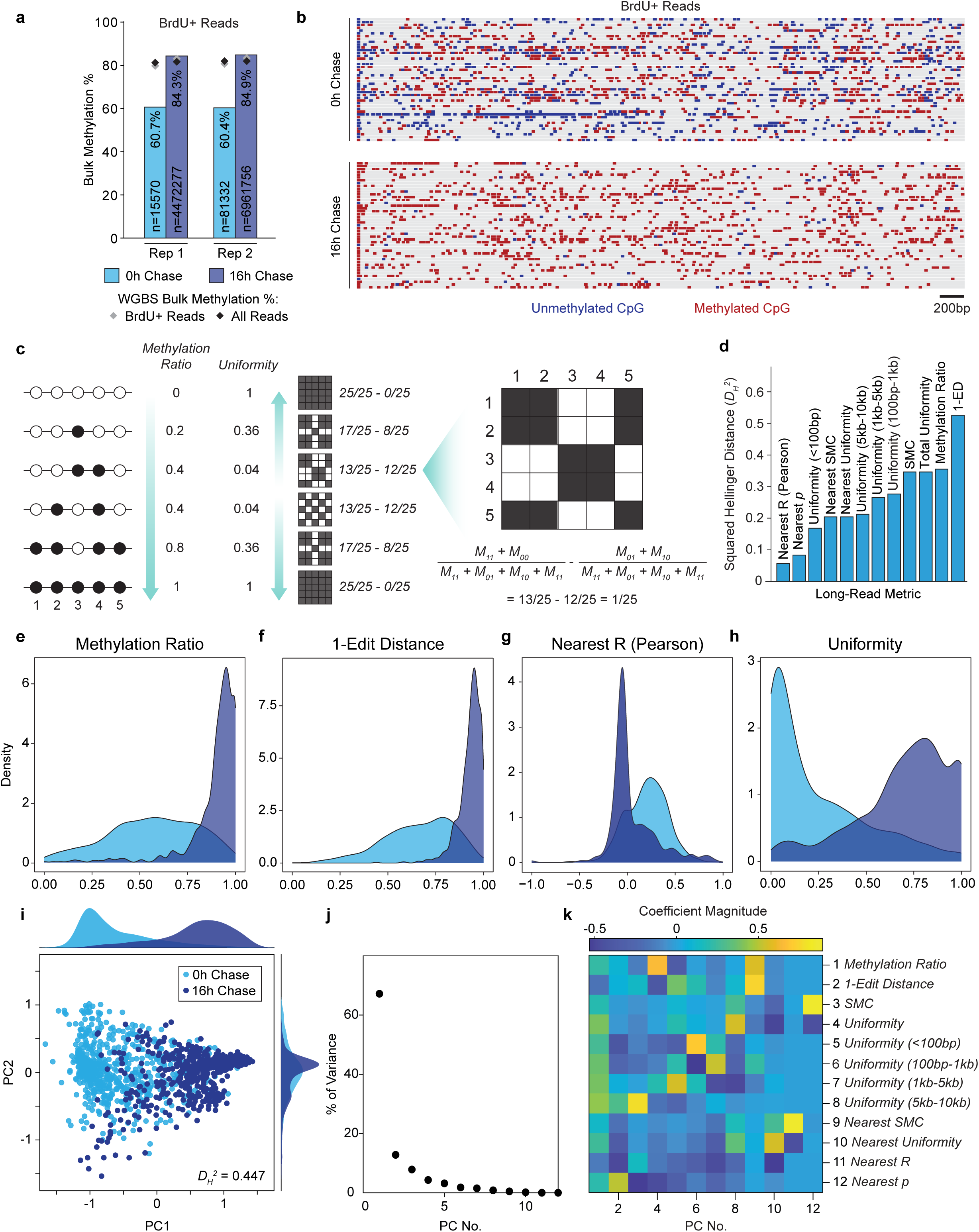
Nascent and mature chromatin can be distinguished by intramolecular CpG methylation patterns. **a)** Bar graph depicting bulk methylation percentage of BrdU-positive reads from BrdU-labeled experimental conditions, as calculated by total methylated CpGs divided by total CpGs. Gray diamond represents the bulk methylation percentage for reads generated by WGBS that capture the corresponding CpGs in BrdU-positive long reads from a given condition. Black diamond represents the bulk methylation percentage for reads generated by WGBS that capture the corresponding CpGs in all long reads from a given condition. **b)** Distributions of CpG methylation across individual BrdU-containing reads in the 0h and 16h chase conditions. Reads are not shown aligned to the genome. **c)** Illustrative schematic depicting the differences in how CpG methylation ratio and uniformity are calculated. CpG sites within reads are represented as circles (black: methylated, white: unmethylated). Matrices represent CpG pairs that are correlated in methylation status (black tiles) or uncorrelated in methylation status (white tiles). **d)** Distributions of methylation ratio for reads derived from the 0h (n = 848) and 16h (n = 975) chase timepoints. **e)** Distributions of 1-edit distance for reads derived from the 0h (n = 848) and 16h (n = 975) chase timepoints. **f)** Distributions of nearest-neighbor Pearson correlation R for reads derived from the 0h (n = 836) and 16h (n = 822) chase timepoints. **g)** Distributions of uniformity for reads derived from the 0h (n = 848) and 16h (n = 975) chase timepoints. For **d-g**, light blue density plot corresponds to 0h chase and dark blue density plot corresponds to 16h chase. **h)** Bar plot depicting the squared Hellinger distance between 0h and 16h read distributions corresponding to each given metric. **i)** PCA based on intramolecular CpG methylation patterns across individual reads (dots). Density plots across PC1 and PC2 are also shown. **j)** Total amount of variance that can be explained by each principal component. **k)** Heatmap representing the coefficients associated with each metric in a given principal component’s corresponding linear combination.

### Nascent and mature chromatin can be distinguished by intramolecular DNA methylation patterns

Upon confirmation that BrdU-labeled nascent chromatin presented an increased proportion of unmethylated CpG sites relative to mature chromatin at both bulk and single-molecule resolution (Fig. 2a,b), we hypothesized that nascent and mature chromatin can be distinguished by intramolecular CpG methylation patterns alone. We sought to devise read-level metrics that could capture not only the proportion of methylated CpGs in a given read, but also other features of CpG methylation patterns related to intramolecular heterogeneity. We first computed read-level metrics previously used for quantitatively measuring epigenetic (methylation ratio and nearest-neighbor Pearson correlation) or genetic (edit distance) differences between DNA molecules, in addition to previously unexplored uniformity metrics (Fig. 2c, Supplemental Note 1) (Methods). The degree of separation achieved by each read-level metric was quantified using a squared Hellinger distance, *D_H_^2^* (Methods) with edit distance having the highest *D_H_^2^* value (Fig. 2d). The distributions of read-level methylation between 0 and 16h reads suggested that mature chromatin is highly methylated, whereas nascent chromatin exhibits widespread variability in methylation ratios (Fig. 2e). To improve the separation between our nascent and mature read populations, we developed a read-level edit distance metric (which computes single nucleotide edit operations required to convert one sequence string to another^20^) that reports how similar the methylation distribution of an individual read is to the corresponding methylation distribution in bulk WGBS data. Edit distance appeared to reduce long-tailed variability in mature reads when compared to distributions of the methylation ratio (Fig. 2f). Previous studies have used Pearson correlations between nearest-neighbor CpG methylation states to quantify single-molecule methylation heterogeneity^13^; however, this metric only modestly separated nascent and mature reads in our data (Fig. 2g, Supplemental Fig. 2a).

To further describe read-level heterogeneity in CpG methylation across single molecules, we used a simple matching coefficient (SMC) to calculate the proportion of matched CpG pairs in each read (Supplemental Fig. 2b). Uniformity metrics were then derived from the SMC to compare the proportions of matched and unmatched CpG pairs within individual reads (Fig. 2c,h). In previous work, we reported that neighboring CpG sites exhibited correlated and faster remethylation kinetics, which could be attributed to enzymatic processivity of DNA methyltransferase 1 (DNMT1)^11^. To address the possibility that enzymatic processivity could impact intramolecular CpG methylation patterns associated with nascent DNA timescales, we also adapted these metrics to account for genomic distance between CpG pairs and methylation states of nearest neighbors (Supplemental Fig. 2c-h).

Toward pseudotime reconstruction of epigenetic modifications across replication-associated timescales, we used principal component analysis (PCA) to cluster our labeled reads according to read-level heterogeneity in CpG methylation, as quantified by linear combinations of the various read-level metrics (Fig. 2i and Supplemental Table 2). Together, our first and second principal components explained ∼79% of variance within the data, with substantial separation between 0 and 16h chase reads, as indicated by a squared Hellinger distance of *D_H_^2^* = 0.447 associated with principal component 1 (PC1) (Fig. 2i,j). Notably, PCA coefficients suggested most metrics were weighted similarly in the calculation of PC1 (Fig. 2k).

To determine if PC1 is more effective at predicting chromatin maturity than other individual read-level metrics, we divided the distributions of 0h and 16h chase reads into 10 equal bins across PC1, as well as each of the 3 read-level metrics with the highest *D_H_^2^* values: Uniformity, Methylation Ratio, and 1-Edit Distance (Supplemental Fig. 2i-l, Supplemental Table 3). We then calculated the proportion of reads in each bin that were derived from either the 0h or 16h chase timepoints and found that, compared to PC1, 1-Edit Distance showed an overrepresentation of 0h chase reads across a majority of the bins, including bins with higher 1-Edit Distance values (Supplemental Fig. 2j,l). This suggested that edit distance is less reliable for distinguishing mature chromatin and supports the use of PC1 as a tool for pseudotime reconstruction. Notably, a subpopulation of reads in our PCA that exhibited lower PC2 values did not appear to be temporally dynamic (according to separation by PC1) (Fig. 2i). This subpopulation appeared to be characterized by nearest-neighbor and short-range intramolecular CpG methylation patterns (Fig. 2k), suggesting that temporal ordering of regions with said features using MPATH could be challenging.

### MPATH recapitulates post-replication CpG methylation dynamics in unlabeled DNA

To investigate the genomic context of reads characterized by low PC2 values, we performed an odds ratio analysis using H1 hESC genomic features annotated via chromHMM (Fig. 3a,b, Supplemental Fig. 3a-b, Supplemental Table 4)^21^. We found that reads with low PC2 values enriched for genomic regions that exhibited hESC-specific hypomethylation, such as promoters and enhancers (Fig. 3a,b, Supplemental Fig. 3a-c, Supplemental Table 4). This finding corroborated with previous observations that genomic regions which are actively engaged in hESCs (e.g., hESC-specific transcription factor binding sites) show invariant bulk methylation levels across replication-associated timescales^10^. Therefore, we concluded that the separation of nascent and mature reads could be improved by triaging reads that appear temporally stagnant according to PC1, marked by lower PC2 values. To isolate our temporally dynamic subpopulation of reads and increase the accuracy of identifying nascent and mature reads, we fitted lines defined by linear inequalities based on BrdU-labeled data to maximize the percentages of true-positive reads in the nascent and mature bins (Fig. 3c) (Methods). This filter resulted in a squared Hellinger distance of *D_H_^2^* = 0.482. The nascent and mature read bins exhibited accuracies of 98.7% (i.e., 98.7% of labeled reads in the nascent bin were correctly identified as nascent 0h reads) and 91.9%, respectively (Fig. 3c).

**Figure 3.**
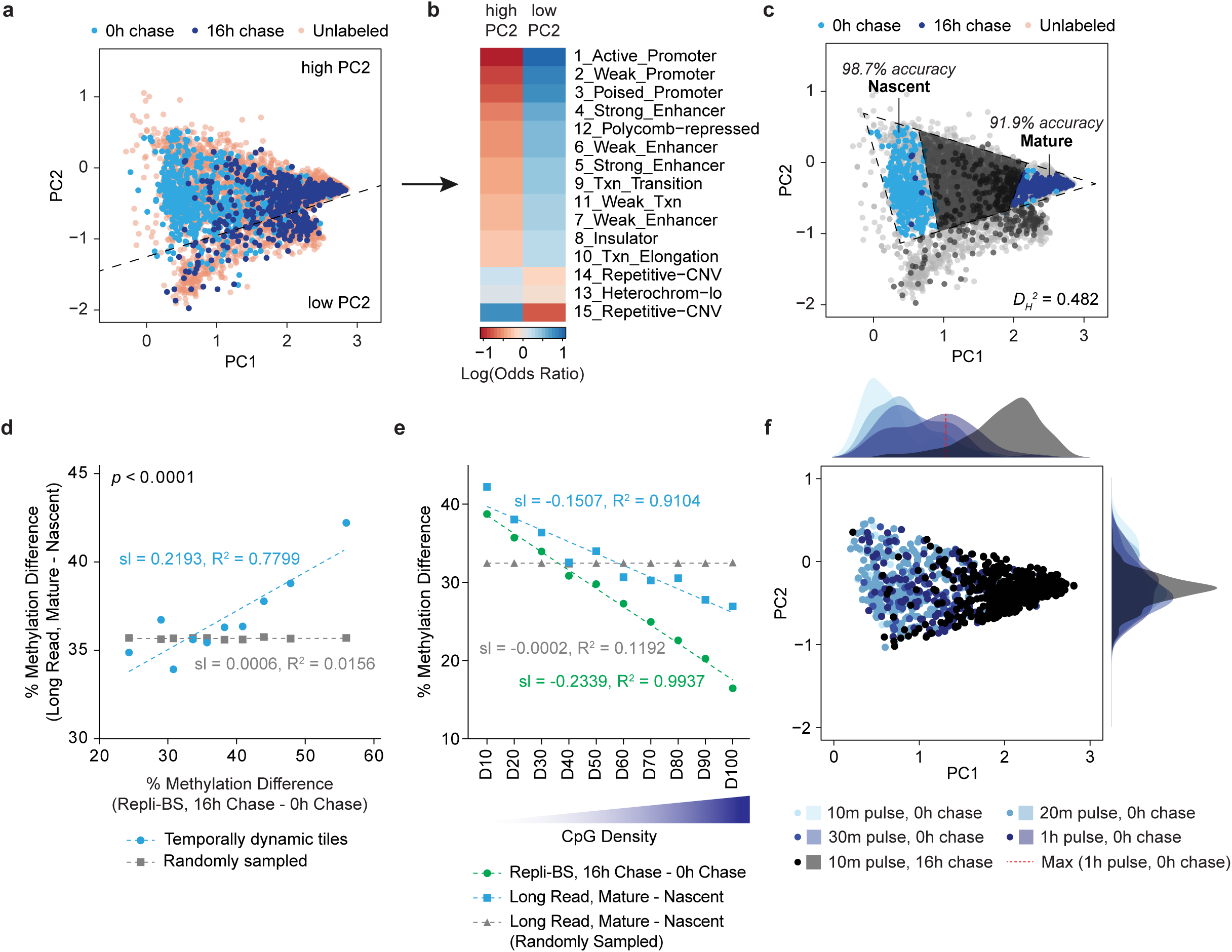
Methylation pseudotime analysis through read-level heterogeneity (MPATH) captures the temporal dynamics of CpG remethylation without DNA labeling. **a)** PCA plot displaying 0h and 16h chase clusters, as well as unlabeled reads (n = 10,000) to which the same linear combinations from the original PCA were applied. Dashed line separates high PC2 and low PC2 read subpopulations. **b)** Heatmap depicting enrichment of high PC2 and low PC2 populations for certain genomic regions annotated by chromHMM. **c)** PCA plot annotated to show the optimized inferred nascent and mature read clusters. **d)** Scatter plot showing the correlation in methylation difference percentage between inferred versus labeled nascent and mature read populations. Dashed lines represent linear regression models applied to the corresponding group. Temporally dynamic tiles were generated from Repli-BS data. The y-axis value corresponding to each temporally dynamic tile was obtained by calculating the bulk CpG methylation percentage of inferred nascent and mature reads that capture the CpGs in a given temporally dynamic tile. For the randomly sampled condition, reads that capture the CpGs in a given temporally dynamic tile were randomly sampled, regardless of inferred post-replication maturity. **e)** Scatter plots and associated linear regression models for methylation difference percentage between labeled versus inferred nascent and mature read populations, as it relates to CpG density. 100 bp tiles were created for each individual CpG in the hg19 reference genome and ranked by CpG density across the tile. **f)** PCA scatter plot depicting the BrdU-containing reads from previous pulse optimization experiments. Linear combinations of read-level metrics associated with the original PC1 and PC2 were applied to the read populations. Density plots across PC1 and PC2 are also shown.

We postulated that the linear combinations of read-level metrics which comprise PC1 and PC2 can be applied to unlabeled reads from cells not treated with BrdU in order to infer nascent and mature read populations. To this end, we sequenced a total of >30 million reads from unlabeled HUES64 hESCs, generating >60x coverage with an N50 read length of 9.3 kb (Supplemental Table 1). We computed read-level metrics for all reads and applied the linear combinations comprising PC1 and PC2 derived from our labeled datasets, followed by triaging of temporally stagnant reads (Fig. 3a,c). We name this pseudotime reconstruction method <u>M</u>ethylation <u>P</u>seudotime <u>A</u>nalysis <u>T</u>hrough read-level <u>H</u>eterogeneity (MPATH). After filtering, MPATH enabled downstream analysis of approximately 9.5 million reads (Supplemental Table 1). To benchmark the ability of MPATH to accurately profile the dynamics of CpG remethylation without DNA labeling, we compared the methylation difference between nascent and mature reads from our unlabeled long-read data set (from which nascent and mature read populations were inferred using MPATH) and publicly available Repli-Bisulfite Sequencing (Repli-BS) data (GSE82045)^10^, in which nascent and mature read populations corresponded to 0 and 16h chase timepoints, respectively. To do this, we first identified differentially methylated regions (DMRs) between 0 and 16h chase data in the Repli-BS study, ordered the DMRs by increasing methylation difference, and grouped them into 10 bins called temporally dynamic tiles, as previously described^22^. This gave us a set of tiles that were ordered according to the magnitude of their time delay in CpG remethylation. We then computed the methylation difference between the corresponding tiled regions captured in our inferred nascent and mature long reads. We observed a high correlation between the methylation differences calculated from inferred versus BrdU-labeled reads, as indicated by an R-squared value of 0.7799 (Fig. 3d), highlighting the ability of MPATH to recapitulate post-replication CpG remethylation dynamics in the absence of DNA labeling.

To investigate the ability of MPATH to capture previously observed CpG density-dependent trends in methylation dynamics, we ranked CpGs according to their local density (+/- 48 bp) and calculated the local methylation difference between inferred nascent and mature reads (Methods). In concordance with previous observations that neighboring CpG sites exhibit faster remethylation kinetics, we found that the methylation difference between nascent and mature reads was inversely correlated with CpG density in both inferred and Repli-BS datasets (Fig. 3e)^10,11^.

We then postulated that PC1 does not only separate nascent and mature read populations but also represents a continuum associated with chromatin maturity, from which a more explicit pseudotime reconstruction could be derived. To test this hypothesis, we applied the linear combinations of read-level metrics associated with PC1 and PC2 to long-read sequences generated from previous BrdU pulse optimization experiments (Fig. 1e,f and Fig. 3f). We observed that the 0h chase reads obtained from various BrdU pulse durations exhibited a positive correlation between BrdU pulse duration and PC1 value (i.e., 0h chase reads appeared more mature with increasing BrdU pulse time), thus confirming the capability of MPATH to temporally order individual long reads across continuous replication-associated timescales (Fig. 3f).

### MPATH enables simultaneous ordering of CpG methylation and hydroxymethylation

Methylated CpGs (5mCG) can be oxidized to hydroxymethylated CpGs (5hmCG) by ten-eleven translocation (TET) enzymes, which leads to active DNA demethylation through base excision repair^23–26^. Given that methylation states can vary during differentiation and in disease, the dynamics of methylation/demethylation turnover have been extensively studied across cell state transitions (e.g., over the course of embryonic development, differentiation, and post-enzymatic deletion)^27–29^. Knockouts of epigenetic writers DNMT3A/B and TETs (5mCG and 5hmCG, respectively) have also suggested that there is active enzymatic competition between DNMTs and TETs at genomic regions termed cDKO-DMRs (differentially methylated regions associated with knockout of *de novo* DNA methyltransferases DNMT3A and 3B)^30^. However, despite previous biochemical analyses suggesting that the removal and production of 5hmCG is replication-dependent^31^, the dynamics of methylation/demethylation turnover during maintenance and throughout the cell cycle are not well understood. Although the post-replication maintenance of 5mCG has been extensively studied, emerging technologies that simultaneously measure 5mCG and 5hmCG have revealed that both modifications are coupled with the cell cycle and generally increase over time^9^, while combinatorial quantification of these marks revealed asymmetry in 5hmCG turnover rates within CpG dyads^32^. While these approaches have enabled deeper insights into the interplay between 5mCG and 5hmCG, the dynamic coordination of these marks across sub-cell-cycle timescales remains unclear.

Furthermore, 5mCG and 5hmCG cannot be distinguished by traditional bisulfite sequencing methods, in which both marks are protected from conversion to uracil^33,34^. While oxidative bisulfite sequencing (oxBS-seq) and enzymatic methyl sequencing (EM-seq) can distinguish between these two base modifications, these technologies still rely on chemical or enzymatic conversion steps that require separate sample preparations^35,36^. In contrast, basecalling algorithms designed for nanopore sequencing can distinguish between 5mCG and 5hmCG through unique electrical traces without the need for conversion, enabling simultaneous profiling of both modifications within a single molecule^37^.

Our pseudotime metrics were calculated based on a 2-state methylation caller, which only distinguishes 5mCG from unmethylated CpGs (CG). To distinguish 5mCG, 5hmCG, and CG marks across pseudotime, we used a 3-state methylation caller. We generated 1-kb tiles across the genome and intersected them with individual long reads from our unlabeled HUES64 dataset to generate 1-kb “single molecules” (Methods). We then calculated the modification ratio and modification uniformity across 10,000 randomly sampled 1-kb single molecules using both methylation callers and ordered them according to pseudotime (Fig. 4a,b) (Methods). While the genome-wide levels of 5mCG and 5hmCG appear to increase following replication, 5hmCG increased to only a fraction of the level observed for 5mCG (0.209 according to the 2-state caller and 0.264 according to the 3-state caller) (Fig. 4a). Interestingly, we found that the 2-state methylation caller overestimates 5mCG modification ratios relative to the 3-state methylation caller (which accounts for the presence of 5hmCG), perhaps due to the 2-state caller’s inability to distinguish 5hmCG from 5mCG. As expected, we observed a pronounced rise in 5mCG uniformity across pseudotime when calculated based on the 2-state methylation caller. However, the 3-state methylation caller showed a thwarted rise in 5mCG uniformity, while 5hmCG exhibited a rapid loss in uniformity (Fig. 4b). Notably, this observation suggests that the deposition of 5hmCG is intimately interspersed among 5mCG marks. Thus, while the propagation of 5mCG is believed to occur *en bloc* via neighbor-guided mechanisms^11,38–40^, conversion of these marks into 5hmCG appeared to occur sporadically throughout the genome during DNA maturation.

**Figure 4.**
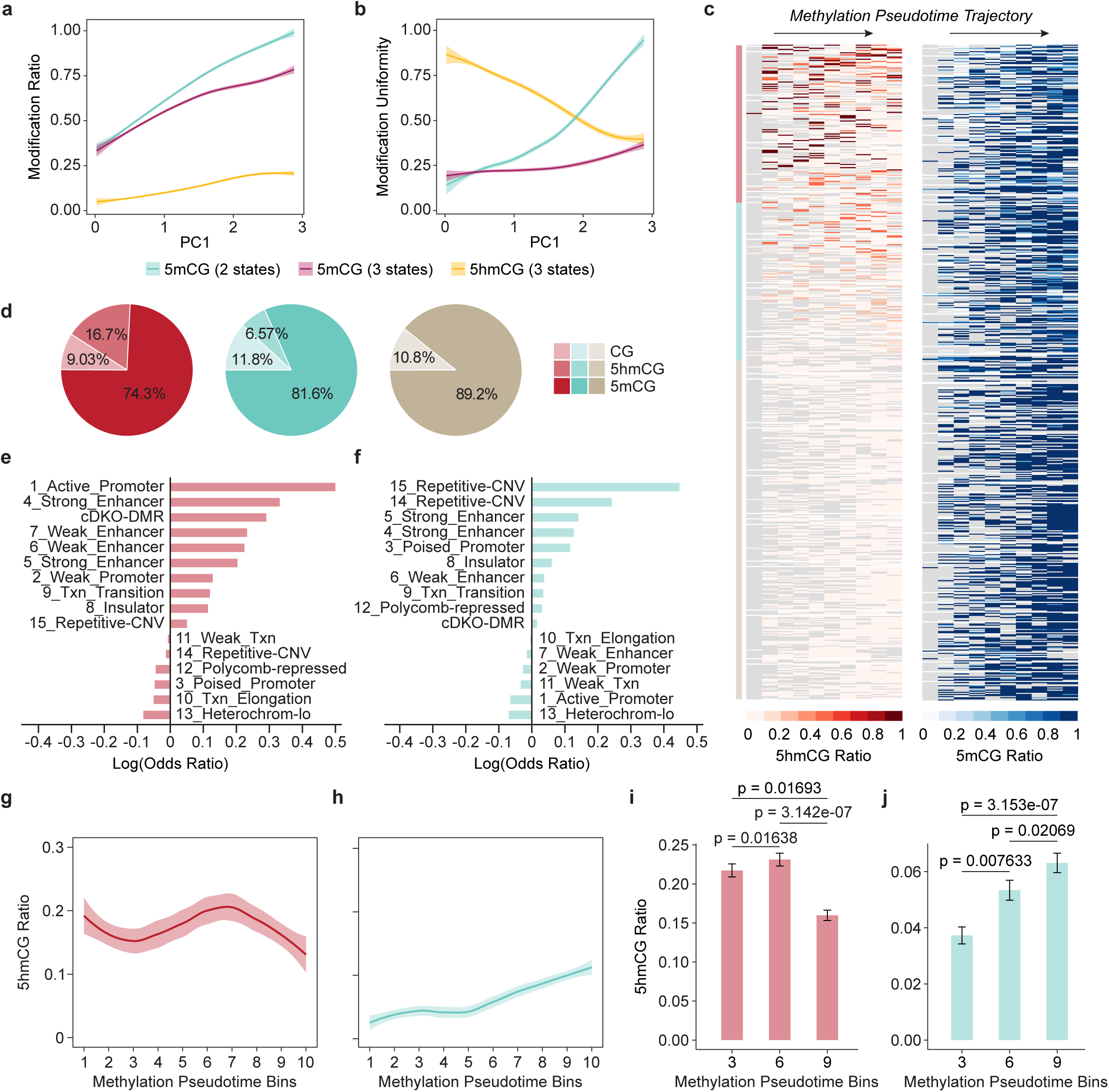
MPATH enables simultaneous profiling of CpG methylation and hydroxymethylation across replication-associated timescales. **a)** Loess smooth curves of CpG methylation ratio and CpG hydroxymethylation ratio, which were calculated for 10,000 randomly sampled 1 kb single molecules using 2 state and 3 state methylation callers. PC1 values for each 1 kb single molecule corresponds to the PC1 value of the read from which the 1 kb single molecule was derived. **b)** Loess smooth curves of CpG methylation uniformity and CpG hydroxymethylation uniformity, which were calculated for 10,000 randomly sampled 1 kb single molecules using 2 state and 3 state methylation callers. PC1 values for each 1 kb single molecule corresponds to the PC1 value of the read from which the 1 kb single molecule was derived. **c)** Heatmaps displaying the CpG methylation ratio and hydroxymethylation ratio for individual CpGs (n = 36,234) across 10 pseudotime bins. Each row represents an individual CpG that is captured at least 10x across pseudotime. CpGs were ordered according to their total hydroxymethylation level across pseudotime and positions are retained between both heatmaps. Red, teal, and beige bars present 3 distinct clusters of CpGs that exhibit high hydroxymethylation, intermediate hydroxymethylation, and no hydroxymethylation across pseudotime, respectively. **d)** Proportion of CpGs that are methylated, hydroxymethylated, and unmethylated in each cluster of the heatmap. **e)** Bar graph depicting enrichment of the highly hydroxymethylated CpG cluster for cDKO-DMRs and certain genomic regions annotated by chromHMM (n = 8,718 CpGs). **f)**. Bar graph depicting enrichment of the intermediately hydroxymethylated CpG cluster for cDKO-DMRs and certain genomic regions annotated by chromHMM (n = 8,718 CpGs). **g-h)** Loess smooth curves of hydroxymethylation ratio of CpGs from the **g)** highly hydroxymethylated CpG cluster (n = 8,718) and **h)** intermediately hydroxymethylated CpG cluster (n = 8,718) across each of the 10 pseudotime bins depicted in panel **c**. **i-j)** Bar plots displaying the hydroxymethylation ratio of CpGs that are captured in pseudotime bins 3, 6, and 9 from the **i)** highly hydroxymethylated CpG cluster (n = 2,120) and **j)** intermediately hydroxymethylated CpG cluster (n = 1,577). Error bars represent standard error and p values are derived from a two-sided Mann-Whitney U test.

### MPATH identifies a subset of CpGs that alternate between methylation states at sub-cell-cycle timescales

Given that 5hmCG turnover has been shown to be cell-cycle-dependent through bulk biochemical analyses in mouse embryonic stem cells^31^, we posited that the base-pair, single-molecule, and temporal resolution provided by MPATH would enable us to investigate methylation/demethylation turnover throughout sub-cell-cycle timescales in hESCs in a locus-specific manner. We filtered for CpGs with at least 10x coverage in our unlabeled dataset (36,234 CpGs or ∼11% of total CpGs captured). Each captured CpG was assigned a PC1 value based on the read it was derived from. We then grouped CpGs into 10 pseudotime bins spanning all PC1 values. We calculated the proportion of CpGs methylated (5mCG ratio) or hydroxymethylated (5hmCG ratio) in each pseudotime bin and clustered CpGs based on their magnitude of hydroxymethylation across all pseudotime bins (Fig. 4c,d) (Methods). We observed that approximately half (51.8%) of the CpG sites never experienced hydroxymethylation across pseudotime. However, of the 17,437 CpGs that did experience hydroxymethylation, we identified two subpopulations that appeared to be either highly or intermediately hydroxymethylated across time (Fig. 4c,d). Interestingly, for highly hydroxymethylated CpGs, methylation and hydroxymethylation marks co-emerge across the majority of pseudotime (Fig. 4c). Hydroxymethylated CpGs, in general, enriched for cis-regulatory elements and repetitive regions, which corroborates their previously reported genomic locations^30,41^. Notably, highly hydroxymethylated CpGs enriched specifically for promoters and enhancers, as well as cDKO-DMRs, which were previously identified as sites of DNMT3A/B and TET competition (Fig. 4e, Supplemental Table 5)^30^. In contrast, intermediately hydroxymethylated CpGs enriched for repetitive regions (Fig. 4f, Supplemental Table 5).

To compare the global dynamics of hydroxymethylation across pseudotime between the two hydroxymethylated clusters, we plotted bulk hydroxymethylation ratios for each pseudotime bin and found that while the intermediately hydroxymethylated cluster (teal) exhibited a gradual accumulation of CpG hydroxymethylation across time, the highly hydroxymethylated cluster (red) did not, suggesting that highly hydroxymethylated CpGs may fluctuate in hydroxymethylation levels during DNA maturation (Fig. 4g,h). Analysis of highly hydroxymethylated CpGs at discrete pseudotime bins 3, 6, and 9 (where only CpGs captured across all three bins were considered) revealed a statistically significant increase, followed by decrease in bulk 5hmCG ratio (Fig. 4i). This was in stark contrast to the monotonically increasing levels of 5hmCG observed in intermediately hydroxymethylated CpGs (Fig. 4j). Thus, these observations revealed a subset of CpGs with greater 5hmCG levels, which were enriched for promoters and enhancers, that actively transition between methylation states at sub-cell-cycle timescales.

### Extension of MPATH to existing NOMe-seq data enables the temporal ordering of chromatin accessibility

The propagation of CpG methylation is often associated with nucleosome occupancy and chromatin condensation^42–44^. However, the relationships between these epigenetic features throughout mitotic inheritance are not well understood. To track chromatin accessibility across sub-cell-cycle timescales and investigate the dynamic coordination between CpG methylation and nucleosome occupancy, we applied MPATH to publicly available long-read NOMe-seq data targeting the *H19/IGF2* imprinted locus in H9 hESCs (with >100x coverage) (GSE183760)^45^. Because the NOMe-seq long reads ranged from 73-84 kb, we created 10 kb sliding windows (with 50% overlap) across the targeted locus, which enabled us to fragment the long reads and apply MPATH to a total of 1,934 read fragments (excluding the imprinting control region, or ICR), which were more similar in length to our labeled reads from which MPATH was established (Fig. 5a). We binned the 10 kb read fragments according to their inferred chromatin maturity, which resulted in a total of 4 read subpopulations (i.e., PC1 bins) ranging from nascent reads (T1 bin) to mature reads (T4 bin) (Methods and Fig. 5a).

**Figure 5.**
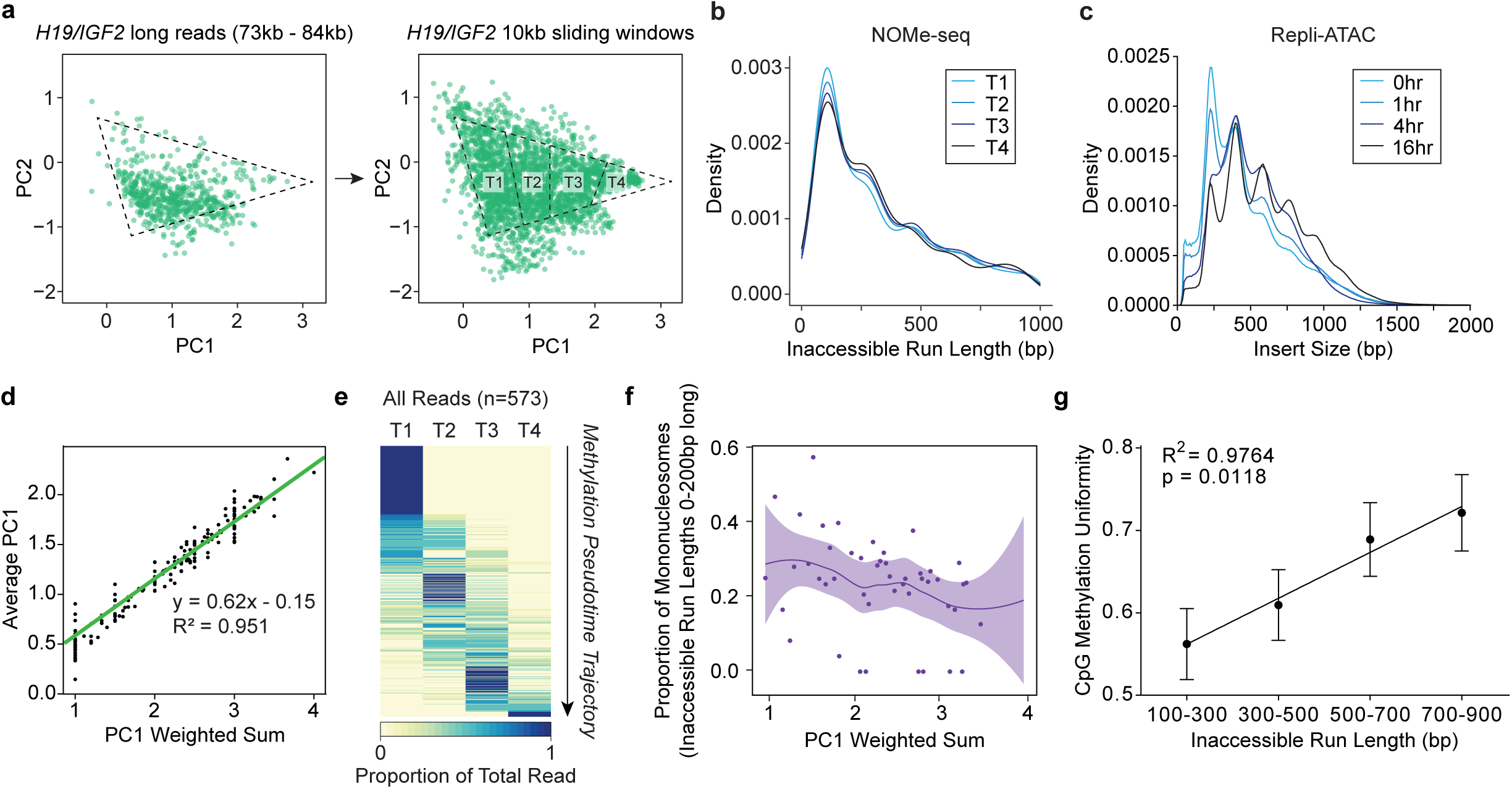
Coupling MPATH with NOMe-seq enables temporal ordering of chromatin accessibility. **a)** Scatter plots representing the original long reads from GSE183760 (left) and individual 10 kb fragments derived from those long reads (right). Each dot represents an individual read of 10 kb fragment. Reads and fragments were transformed using the linear combinations comprising the original PC1 and PC2. PC1 was divided into 4 bins T1-T4, ordered from most nascent to most mature. **b)** Density plots showing the distributions of inaccessible run lengths for 10 kb fragments that fall into each of the PC1 bins (T1-T4). T1, n = 7,866. T2, n = 5,550. T3, n = 5,810. T4, n = 2,402. **c)** Density plots showing the distributions of insert size associated with each chase timepoint in Repli-ATAC. 0hr, n = 10,863,954. 1hr, n = 22,901,076. 4hr, n = 37,219,193. 16hr, n = 40,564,395. **d)** Scatter plot and linear regression model depicting the correlation between an average of PC1 values from individual 10 kb fragments comprising a long read versus a weighted sum of PC1 bins for individual 10 kb fragments comprising a long read. **e)** Heatmap depicting the proportion of 10 kb fragments comprising an individual long read that fall into each PC1 bin. Each row represents an individual long read. Reads are ordered by their weighted sum of PC1 bins. **f)** Scatter plot and Loess smooth curve depicting the proportion of inaccessible run lengths from individual 10 kb fragments that are 0-200 bp long, plotted across the PC1 weighted sum of the 10 kb fragment. Only 10 kb fragments from Cluster 5 are shown. **g)** Scatter plot and linear regression model depicting the correlation between inaccessible run length and CpG methylation uniformity across 10 kb fragments from Cluster 5. R^2^ was calculated from mean CpG methylation uniformity values. Error bars represent standard error. p value was derived from an F-test.

Next, we sought to utilize MPATH to temporally profile chromatin accessibility (measured through exogenous GpC methylation) at the *H19/IGF2* locus within the NOMe-seq data set. We identified regions within read fragments that were accessible (GpC methylated) and inaccessible (GpC unmethylated) to the M.CviPI GpC methyltransferase and observed that the distribution of inaccessible runs was characterized by run lengths reminiscent of nucleosomes (Fig. 5b). For example, the first peak occurs at a run length of approximately 130 bp, which is reminiscent of the length of DNA wrapped around a mononucleosome (147 bp), whereas successive peaks seem indicative of closely packed dinucleosomes, trinucleosomes, etc. Interestingly, separating the read fragments according to PC1 bin revealed that T1 (nascent) read fragments were most enriched for mononucleosomes (Fig. 5b). This result supported previous observations that nascent chromatin is generally accessible and gradually condenses as the epigenetic landscape is restored^46^. Accordingly, T4 (mature) read fragments were least enriched for mononucleosomes. To validate the results obtained from coupling MPATH with NOMe-seq, we performed a replication-coupled assay for transposase-accessible chromatin followed by sequencing (Repli-ATAC sequencing), which applies mitotic labeling to map chromatin accessibility across replication-associated timescales (Supplemental Fig. 4a)^47^. The distributions of insert size presented similar trends (i.e., the enrichment of mononucleosomes within nascent reads from the 0h chase timepoint) (Fig. 5c). We also found that MPATH could identify temporal heterogeneity within sub-nucleosomal chromatin accessibility patterns, which could potentially be attributed to the dynamics of various chromatin-associated factors (including transcription factors) throughout replication (Supplemental Fig. 4b).

Binning of read fragments also enabled us to investigate if various sites across a single DNA molecule can exhibit different chromatin maturation dynamics in their local epigenetic landscape. To this end, we calculated a weighted sum of PC1 bins for each individual read, defined as the sum of the proportions of read fragments assigned to each weighted PC1 bin, otherwise referred to as a PC1 weighted sum (Supplemental Note 1). We show that the PC1 weighted sum is directly proportional to the average PC1 across all 10 kb fragments of an individual long read and clustered the long reads according to this metric (Fig. 5d,e). Interestingly, we found that while 237 long reads (or 41.4% of total long reads) were composed of read fragments derived from only one PC1 bin, 336 long reads (58.6%) exhibited variation in PC1 values assigned to their corresponding read fragments, suggesting that during chromatin maturation, temporal heterogeneity can exist in single molecules across large genomic regions (Fig. 5e).

To investigate the coordinated dynamics of nucleosome occupancy and CpG methylation across sub-cell-cycle timescales, we first isolated NOMe-seq long reads generated from a 73 kb targeted region within the *H19/IGF2* locus^45^ and clustered captured GpCs according to their read coverage to ensure fair evaluations of locus-specific trends in chromatin accessibility (Supplemental Fig. 4c-e). For Cluster 5, which encompasses the ICR, we observed a decrease in short inaccessible run lengths reminiscent of mononucleosomes throughout pseudotime (Fig. 5f, Supplemental Fig. 4f-j). Increasing chromatin inaccessibility in this cluster also appeared to correlate with increasing CpG methylation uniformity, suggesting that nucleosome compaction and CpG methylation uniformity are coupled across sub-cell-cycle timescales (Fig. 5g).

### Phasing of long-read nanopore data reveals allele-specific trends in pseudotime distribution associated with X chromosome activity

Previous studies have suggested that alleles can exhibit different dynamics during genome replication^48–50^. In the context of X chromosome inactivation (XCI), the inactive X chromosome (X_i_) undergoes rapid and uniform replication during late S-phase of the cell cycle^51,52^. Thus, we posited that MPATH could discern X chromosome activity based on allelic differences in pseudotime distribution. We sequenced genomic DNA from Abl.1 mouse B cells to >40x coverage (N50 read length: 16.1 kb) and phased reads at a given 1 Mbp locus on the X chromosome (chrX:103,530,980-104,531,001) into two haplotypes (Fig. 6a, Methods). We determined that haplotype 1 was derived from X_i_ due to the unmethylated state of the *Xist* promoter (Fig. 6a). We then created 20 kb windows across the locus and identified windows at which both haplotypes exhibited similar bulk CpG methylation levels (<15% bulk CpG methylation difference) to mitigate the influence of XCI-associated methylation differences on pseudotime reconstruction (Fig. 6b). Then, we applied MPATH to all reads within these windows and found that reads derived from X_i_ exhibited a more uniform distribution of PC1, which could reflect more uniform (and rapid) chromatin maturation kinetics associated with X_i_ (Fig. 6c)^51^. In contrast, phased reads derived from an autosome (chr10) exhibited very similar distributions of PC1, thus suggesting that alleles can progress differently across replication-associated timescales in the case of monoallelic expression (Fig. 6d).

**Figure 6.**
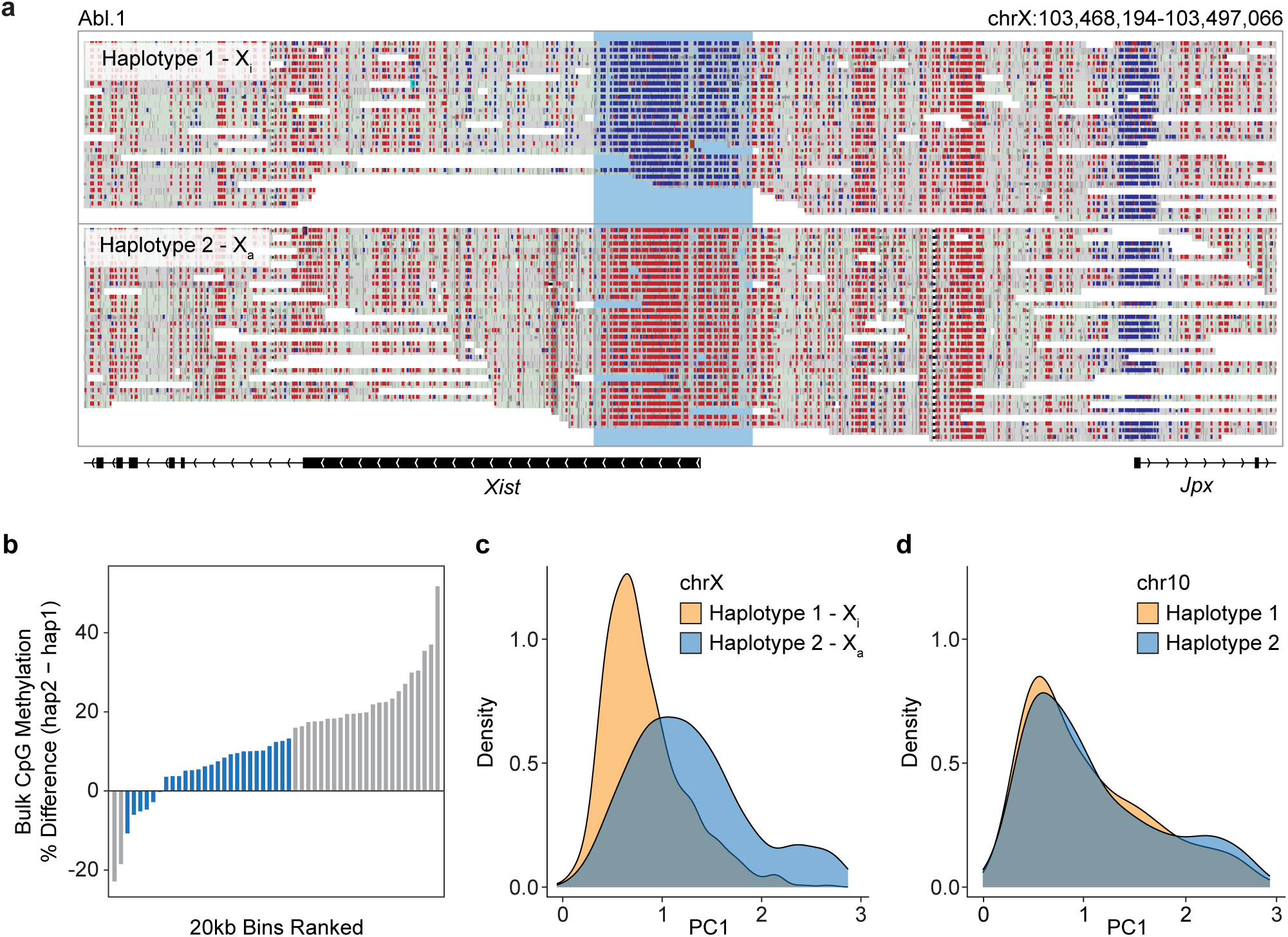
MPATH reveals allele-specific trends in pseudotime distribution associated with genomic imprinting and X chromosome inactivation. **a)** Browser window depicting phased long reads from Abl.1 mouse B cells across chrX:103,468,194-103,497,066. Colored according to methylation status of CpG sites (blue: unmethylated, red: methylated). Haplotype 1 is derived from the inactive X, as shown through lack of CpG methylation at the *Xist* promoter (shaded blue region). **b)** Bar plot depicting the difference in bulk CpG methylation between reads derived from haplotype 1 vs 2 across 51 20-kb windows. Bars that are blue represent windows at which the difference in bulk CpG methylation between reads derived from haplotype 1 vs 2 is ≤ 15%. **c)** Density plot depicting the distribution of PC1 values assigned to reads derived from haplotype 1 (n = 637) vs haplotype 2 (n = 600) within filtered windows across the X chromosome region (chrX: 103,530,980-104,531,001) at which the difference in bulk CpG methylation between reads derived from haplotype 1 vs 2 is ≤ 15%. **d)** Density plot depicting the distribution of PC1 values assigned to downsampled reads derived from haplotype 1 (n = 637) vs haplotype 2 (n = 600) within a region of the autosome chr10.

## Discussion

In this study, we presented a novel label- and conversion-free method that leverages intramolecular CpG methylation patterns for methylation pseudotime analysis of single DNA molecules. We demonstrated that when coupled with DNA modification mapping strategies, MPATH can be particularly valuable for temporally ordering epigenetic modifications without the need for cytotoxic nucleoside analogs or chemical conversion steps. Although actively engaged regions exhibit stagnant mean methylation levels throughout the cell cycle^10^, applying MPATH to long-read sequences enabled the temporal ordering of these elements using surrounding regions within individual reads. Coupling MPATH with a 3-state methylation caller enabled unprecedented temporal resolution into DNA methylation/demethylation turnover during stem cell maintenance, revealing a subset of CpGs which alternate between hydroxymethylated and unhydroxymethylated states at sub-cell-cycle timescales within cis-regulatory elements, potentially reflecting competition between DNA methylation writer and eraser enzymes^28,30,53–55^. We also showed that retroactive application of MPATH could provide temporal context to an existing long-read NOMe-seq dataset, revealing patterns of CpG methylation and nucleosome occupancy that were coordinated in time. These findings support the potential of MPATH as a framework for investigating the extent to which these coordinated patterns are shaped through post-replication epigenetic crosstalk. Finally, extension of MPATH to phased reads revealed how alleles can progress differently through replication-associated timescales according to X chromosome activity. Therefore, by providing temporal context to phased long-read datasets, MPATH offers a novel approach for studying the mechanistic underpinnings of monoallelic gene expression.

MPATH utilizes linear combinations of discrete read-level metrics generated based on PCA to infer chromatin maturity. In contrast to other trajectory inference methods that utilize machine learning or dynamical modeling^56–59^, the advantages of our approach include the ability to benchmark our pseudotime scores against “ground truth” labeled data and identify specific features of methylation patterning that drive pseudotime reconstruction. Many multimodal sequencing strategies have already been adapted to long-read sequencing platforms, which will support diverse applications of MPATH^60–62^. Future efforts will benefit from extending MPATH to other cell types and synthetic cellular systems. We posit that MPATH will be especially useful for deciphering epigenetic crosstalk and understanding chromatin dynamics during replication and cell state transitions at sub-cell-cycle time resolution.

## Methods

### Cell culture

HUES64 hESCs were cultured on Geltrex (Thermo Fisher Scientific, A1413302) coated tissue culture plates with mTeSR (STEMCELL Technologies) media. Cells were incubated at 37°C with 5% CO_2_. Media was changed daily and cells were passaged upon reaching 80% confluency using ReleSR (STEMCELL Technologies). v-Abl pro-B clonal cell line Abl.1, derived previously from 129S1/SvImJ × CAST/EiJ F1 female mice^63^, were cultured in RPMI medium (Gibco) containing 15% FBS (Sigma), 1X L-Glutamine (Gibco), 1X Penicillin/Streptomycin (Gibco), and 0.1% β-mercaptoethanol (Sigma).

### Cell cycle arrest

hESCs were treated with 2 µg/ml nocodazole for 16 h, collected, and washed twice with DPBS (Gibco, 14190144) prior to genomic DNA extraction by phenol-chloroform extraction.

### BrdU labeling

To label nascent DNA, 5-bromo-2′-deoxyuridine (BrdU) (Sigma-Aldrich, B5002) was added to the cell culture medium at a final concentration of 50 µM or 500 µM. Cells were incubated in labeled medium for 10 minutes at 37°C with 5% CO_2_ (for BrdU pulse optimization, incubation times of 20, 30, and 60 minutes were also used). Next, nascent samples were washed once with DPBS (Gibco, 14190144), and cells were collected using Accutase (STEMCELL Technologies), pelleted, and fixed in 75% ethanol by adding 100% ethanol dropwise to the cell resuspension until a final ethanol concentration of 75% was reached. In contrast, mature samples were washed three times with DPBS, and further incubated in fresh mTeSR. At 16 hours of incubation, mature samples were collected, pelleted, and fixed in 75% ethanol as previously described.

### Assessing cell viability

Collected cells were stained using the LIVE/DEAD™ Fixable Violet Dead Cell Stain Kit (Thermo Fisher Scientific, L34955) according to the manufacturer’s instructions. Samples were analyzed via flow cytometry using a NovoCyte 3000 (Agilent Technologies).

### Genomic DNA isolation

For samples that were arrested with nocodazole or labeled with 50 µM BrdU, cells were lysed via incubation in SDS-PK buffer (0.5% wt/vol) sodium dodecyl sulfate (SDS), 50 mM Tris-HCl, 0.01 M EDTA, 1 M NaCl and 0.2 mg/ml proteinase K) at 56 °C for 2 h. Genomic DNA was isolated using phenol-chloroform extraction and isopropanol precipitation. For samples labeled with 500 µM BrdU, genomic DNA was isolated using the Puregene Cell Kit (8 x 10^8^) (Qiagen, 158043) or Monarch^®^ Genomic DNA Purification Kit (NEB, T3010S) according to the manufacturer’s instructions. Abl.1 B cells were collected on day 5, and debris were removed using Histopaque. DNA was prepared using Sigma GenElute kit, and then enriched for longer fragments using Circulomics SRE XS kit.

### DNA shearing for nanopore sequencing

For 500 µM BrdU labeled samples and unlabeled samples, genomic DNA was sheared to approximately 6-8 kb using g-TUBEs (Covaris, SKU 520079) by spinning 2-3 μg of unfragmented gDNA at 7,200 rpm for 1 min in a Pico 21 Microcentrifuge (Thermo Scientific). Then, tubes were inverted and spun down under the same conditions.

### Nanopore sequencing

For samples labeled with 50 µM BrdU, libraries were prepared from purified and fragmented genomic DNA using a Ligation Sequencing Kit (Oxford Nanopore Technologies, SQK-LSK110) according to the manufacturer’s instructions. Briefly, DNA repair and end-prep were performed using NEBNext^®^ FFPE DNA Repair Mix (NEB, M6630) and NEBNext^®^ Ultra II End Repair/dA-tailing Module (NEB, E7546). Adapter ligation and clean-up were performed using the NEBNext^®^ Quick Ligation Module (NEB, E6056). Long Fragment Buffer from the Ligation Sequencing Kit was used in the clean-up step to enrich for DNA fragments of >3 kb. Libraries were loaded onto R9.4.1 flow cells (Oxford Nanopore Technologies, FLO-MIN106D) and sequenced using a MinION Mk1b device (Oxford Nanopore Technologies) for 16 hours. Minimum read length was set to 200 bp. For Abl.1 B cells, libraries were prepared similarly, loaded onto R9.4.1 flow cells, and sequenced using a PromethION P2 solo device (Oxford Nanopore Technologies). For samples labeled with 500 µM BrdU and samples not labeled with BrdU, libraries were prepared using a Ligation Sequencing Kit (Oxford Nanopore Technologies, SQK-LSK114) according to the manufacturer’s instructions, similarly to previously described. Libraries for samples labeled with 500 µM BrdU were loaded onto R10.4.1 flow cells (Oxford Nanopore Technologies, FLO-MIN114) and sequenced using a MinION Mk1b device for 32 hours. Libraries generated from unlabeled samples were loaded onto R10.4.1 flow cells (Oxford Nanopore Technologies, FLO-PRO114M) and sequenced on a PromethION P2 solo device (Oxford Nanopore Technologies) for 72 hours. Minimum read length was set to 1000 bp. All sequencing data was collected using MinKNOW v23.07.5.

### Preprocessing of nanopore data

For BrdU-labeled samples loaded on R9.4.1 flow cells (Oxford Nanopore Technologies, FLO-MIN106D), basecalling on raw current signals was performed using Guppy v6.5.7. DNA sequences were aligned to the hg19 human reference genome using Minimap2 v2.24 and chimeric reads were removed using samtools v1.15.1. To identify BrdU-containing reads, DNAscent v3.1.2^64^ was used to detect the BrdU base analog within individual reads. CpG methylation was called using nanopolish v0.13.2^65^. For BrdU-labeled samples loaded on R10.4.1 flow cells (Oxford Nanopore Technologies, FLO-MIN114) and unlabeled samples loaded on R10.4.1 flow cells (Oxford Nanopore Technologies, FLO-PRO114M), basecalling and alignment to the hg19 human reference genome were performed using Dorado v0.6.2. Chimeric reads were removed using samtools v1.15.1. To identify BrdU-containing reads, DNAscent v4.0.1 was used to detect the nucleoside analog within individual reads. CpG methylation was called using Dorado v0.6.2. Nanoplot v1.41.6 was used to assess and visualize read-level statistics such as number of reads, read length N50, mean and median read length, and mean and median read quality. For mouse B cells raw nanopore reads were basecalled and CpG methylation detection was done using Guppy v6.3.9 (basecall_model=2021-05-05_dna_r9.4.1_promethion_384_dd219f32). Reads were aligned to the mm10 mouse reference genome using Minimap2 v2.22 with default parameters.

### Calculating read-level metrics

For each individual read, average BrdU is defined as the average of the probabilities that each thymidine is BrdU, as determined from DNAscent v4.0.1. A mock-treated sample with no BrdU was used to determine a threshold average BrdU value, above which a read is determined to be BrdU-containing. Methylation ratio is defined as the ratio of total methylated CpGs to total CpGs within the read. The uniformity of a read is defined as the proportion of matched CpG pairs minus the proportion of unmatched CpG pairs within the read. Detailed information and equations for all read-level metrics, including metrics that are adapted to account for nearest neighbors or genomic distance, can be found in the Supplemental material (Supplemental Note 1).

### Dimensionality reduction for methylation pseudotime analysis

Read-level metrics were integrated into a principal component analysis (PCA) to enable methylation pseudotime analysis. PCA was conducted using MATLAB (R2023b).

### Applying linear combinations generated by MPATH to unlabeled data

Each principal component in the PCA on labeled data was characterized by a linear combination of the input read-level metrics, where each individual read-level metric was associated with a unique coefficient for each principal component. Read-level metrics were calculated for reads derived from unlabeled data. Then, the coefficients of the linear combinations comprising the principal components derived from labeled data were applied to the read-level metrics calculated from unlabeled data.

### Sourcing genomic annotations

All annotated genomic regions were previously inferred through chromHMM^21^ and downloaded from the UCSC Table Browser^66^. Promoter regions were defined as 2 kb upstream and 500 bp downstream of transcription start sites (TSS). Intergenic regions were determined using the BEDTools “subtract” function^67^. The BEDTools “intersect” function was used to determine which regions were captured by sequenced long reads.

### Enrichment analyses

Odds ratio (OR) analyses were performed to determine the strength of the association of read populations of interest or CpGs of interest with particular genomic regions. For enrichment analysis of low PC2 reads, OR was calculated as 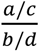, where *a* = number of reads that are categorized as low PC2 and capture the genomic region of interest, *b* = number of reads that are categorized as low PC2 and do not capture the genomic region of interest, *c* = number of reads that are not categorized as low PC2 reads and capture the genomic region of interest, and *d* = number of reads that are not categorized as low PC2 reads and do not capture the genomic region of interest. For enrichment analysis of hydroxymethylation clusters, *a* = number of CpGs within a given cluster that occur within the genomic region of interest, *b* = number of CpGs within a given cluster that occur outside of the genomic region of interest, *c* = number of CpGs that are not within a given cluster that occur within the genomic region of interest, and *d* = number of CpGs that are not within a given cluster that occur outside of the genomic region of interest. The logarithmic OR value (log(OR)) was calculated for each genomic region of interest. The Fisher’s exact test was used to determine the statistical significance of ORs. Heatmaps were created using the R package pheatmap v1.0.12.

### Binning read populations across pseudotime

Three linear inequalities across PC1 and PC2 defined the population of reads that can be analyzed by MPATH (Supplemental Note 1). These linear inequalities were manually optimized to encapsulate reads that are temporally dynamic, as determined by the placement of BrdU-labeled reads. Two linear inequalities were then curated to define nascent and mature reads within this population. The latter linear inequalities were manually optimized to maximize the proportion of true nascent reads in the nascent bin and true mature bins in the mature bin. To create 4 bins across pseudotime, the population of reads that were not categorized as nascent or mature were split into 2 more bins according to the PC1 value associated with the peak of the BrdU 10 minute pulse, 0h chase read distribution across PC1 (Figure 3F).

### Quantifying CpG hydroxymethylation across pseudotime

CpG hydroxymethylation was called on long reads derived from unlabeled hESCs using Dorado v0.6.2, as described above. Each read was assigned a PC1 value using MPATH as described above. To profile CpG hydroxymethylation across pseudotime globally, we generated 1 kb tiles of the hg19 human reference genome and intersected this with long reads in order to create 1 kb single molecules. We calculated the methylation ratio and uniformity across 10,000 randomly sampled 1 kb single molecules (Supplemental Note 1) and plotted these values against the PC1 value corresponding to the read from which the 1 kb single molecule was derived as a smooth curve using the R package ggplot2 v3.5.1. All possible PC1 values from this data set were categorized into 10 equal-sized bins. To profile CpG hydroxymethylation across pseudotime locus-specifically, we first filtered for CpGs with ≥10x coverage. For each of these CpGs, we calculated the proportion of reads within each PC1 bin that capture that CpG in a hydroxymethylated or methylated state. This data was plotted as a heatmap using pheatmap v1.0.12. Clusters of hydroxymethylated CpGs were determined by first categorizing CpGs into CpGs that experience hydroxymethylation vs no hydroxymethylation. Then, the hydroxymethylated cluster was divided evenly into 2 sub-clusters, where the first cluster is considered highly hydroxymethylated and the second cluster is considered intermediately hydroxymethylated.

### Processing publicly available NOMe-seq data

Raw nanopore reads from NOMe-seq conducted on H9 hESCs were downloaded in fast5 format from the National Center for Biotechnology Information (NCBI) under Bioproject ID PRJNA510783. Basecalling, CpG methylation calling, GpC methylation calling, and alignment to the hg19 human reference genome were all performed using Dorado v0.6.2. Chimeric reads were removed using samtools v1.15.1. To avoid ambiguous methylation outputs, CpGs and GpCs in a GCG context were excluded for downstream analysis (26.4% of CpGs and 18.5% of GpCs).

### Inaccessible Run Lengths

We first created 10 kb windows of the hg38 human reference genome and intersected them with the CpG and GpC methylation data from NOMe-seq long reads to create 10 kb single molecules. The start and end coordinates of runs within a molecule were defined as the midpoint between the first or last methylated GpC and nearest unmethylated GpC. Inaccessible runs were characterized by runs in which all GpCs were unmethylated.

### Repli-ATAC sequencing

HUES64 hESCs were grown to ∼70% confluency in a 6-well plate. Different chase timepoints were assigned to different wells. To label nascent DNA, cell culture medium was replaced with 2 mL/well of mTeSR containing 50 µM BrdU. Cells were incubated in labeled medium for 1 hour at 37°C with 5% CO_2_. Next, 0h chase samples were washed once with mTeSR, and cells were collected using Accutase. For later chase timepoints, two washes with mTeSR were done before incubating the cells at 37°C with 5% CO_2_ until the chase timepoint was reached. Then, cells were collected using Accutase. Collected cells were resuspended in 1 mL ice-cold DPBS buffer and 100K cells were transferred to and pelleted in a 1.5 mL Eppendorf tube at 4°C. The pellet was resuspended in 50 uL cold lysis buffer (10 mM Tris-HCl, pH 7.4, 10 mM NaCl, 3 mM MgCl_2_, 0.1% IGEPAL CA-630). Nuclei were spun down immediately at 500 x g for 10 minutes at 4°C, supernatant was discarded, and the pellet was stored on ice.

To initiate the transposition reaction, the nuclei were resuspended in transposition reaction mix: 50 µL TD (2x reaction buffer, Illumina, FC-121-1030), 5 µL TDE1 (Nextera Tn5 Transposase, Illumina, FC-121-1030), and 45 µL nuclease-free water (Invitrogen, 10977-015). The transposition reaction was incubated on a thermomixer at 37°C for 30 minutes. Then, tagmented DNA was purified using the MinElute PCR Purification kit (Qiagen, 28004). To amplify transposed DNA fragments from each sample, the following was combined in a 0.2 mL PCR tube: 20 µL transposed DNA, 5 µL nuclease-free water, 25 µL NEBNext High-Fidelity 2x PCR Master Mix (NEB, M0541). Samples were placed in a thermal cycler for 1 cycle of 72°C for 5 minutes to allow for extension of both ends of the primer after transposition. DNA was purified using the MinElute PCR Purification kit.

10 µL of each sample was used for BrdU immunoprecipitation (IP) and 10 µL was stored aside at 4°C as a non-IP control. To denature the DNA in the IP sample, 490 µL of TE buffer was added and the sample was incubated at 95°C for 5 minutes on a thermomixer before cooling in an ice bath. The denatured DNA was collected and added to 60 µL of 10x IP buffer (28.5 mL nuclease-free water, 5 mL of 1 M sodium phosphate, pH 7.0, 14 mL of 5 M NaCl, and 2.5 mL of 10% wt/vol Triton X-100). 40 µL of 12.5 µg/mL anti-BrdU antibody (BD Biosciences, 555627) was added, and the sample was incubated for 1 hour at room temperature with constant rotation. The sample was centrifuged at 17,000 x g for 5 minutes at 4°C and the supernatant was completely removed. 750 µL of ice-cold 1x IP buffer was added before centrifuging and removing the supernatant again.

The pellet was then resuspended in 200 µL digestion buffer (50 mM Tris-HCl, pH 8.0, 10 mM EDTA, 0.5% SDS) with 0.25 mg/mL Proteinase K and incubated overnight at 37°C, then for 60 minutes at 56°C. Next, the sample was purified using AMPure XP Beads (Beckman Coulter, A63881) (2x bead volume) and resuspended in 20 µL Elution Buffer. To amplify the transposed DNA fragments, the following was combined in a 0.2 mL PCR tube: 20 µL transposed DNA, 2.5 µL 25 µM Custom Nextera PCR Primer 1, 2.5 µL 25 µM Custom Nextera PCR Primer 2, 25 µL NEBNext High-Fidelity 2x PCR Master Mix. The following PCR conditions were used: 1 cycle of 72°C for 5 minutes, 98°C for 30 seconds, followed by 5 cycles of 98°C for 10 seconds, 63°C for 30 seconds, 72°C for 1 minute. The PCR reaction was monitored using qPCR to stop amplification before saturation, in order to reduce GC and size bias^68^. Finally, the amplified library was purified using the MinElute PCR Purification kit.

### Repli-ATAC sequencing data processing

Fastq files were trimmed using Trim Galore v0.4.4^69^ and aligned using BWA^70^. Samtools v1.15.1 was used to remove mitochondrial DNA, duplicate reads, and unmapped reads. BamTools^71^ was used to remove reads containing more than 4 mismatches and reads with insert size greater than 2 kb. Peak calling was performed using MACS2^72^. To quantify insert sizes, the *CollectInsertSizeMetrics* function of Picard^73^ was used with M=0.5 for each bam file. Histogram counts were merged across 2 replicates and plotted as a density of all reads as normalization.

### Phasing long-read data

To phase the long reads from the publicly available NOMe-seq data set (GSE183760), reads that capture the imprinting control region (ICR) at the *H19/IGF2* locus were assigned an MPATH score according to their intramolecular CpG methylation patterns, excluding the CpGs within the ICR. To determine if a read was derived from the maternal or paternal allele, we first calculated the fraction of methylated CpGs in the ICR for each read. The ICR was considered to be methylated if the fraction of methylated CpGs was >0.5 and unmethylated if the fraction of methylated CpGs was <0.5. Reads with methylated ICRs were assigned to the paternal allele while reads with unmethylated ICRs were assigned to the maternal allele, in corroboration with the known methylation states of the paternal and maternal ICRs for the *H19/IGF2* locus. Genomic variants for the 129S1/SvImJ and CAST/EiJ mouse strains were obtained from The Jackson Laboratory database. Homozygous SNP sites from each strain were used to construct a VCF file representing the F1 cross. Read phasing was performed using Whatshap v2.0.

### Statistical analyses

Statistical analyses were performed using GraphPad Prism v9.3.1 for Windows (linear regression models) and in R v4.0.3 (Fisher’s exact test).

## Data availability

Raw sequencing data and processed data for this study are available through the Gene Expression Omnibus. Data associated with Figures 1-5 can be accessed at GSE290978 (Nanopore) and GSE290974 (ATAC-seq). Raw data in POD5 or FAST5 format for Figures 1-5 are available upon request. Both raw and processed data associated with Figure 6 can be accessed at GSE286139.

## Code availability

Source code for analysis is available at https://github.com/downinglab/mpath/.

## Acknowledgements

We thank M. Byun for insightful discussions and M. Boemo for sharing a pre-release version of DNAscent that helped toward preliminary analyses for this study. This work was partially supported by NSF grants (EF2022182 and DMS1763272), a grant from the Simons Foundation (594598, QN), and an NIH New Innovator Award (DP2) Grant DP2CA250382-01 to T.L.D. A.T. was supported by a training grant from the California Institute for Regenerative Medicine under Award Number EDUC4-12822. The content is solely the responsibility of the authors and does not necessarily represent the official views of the California Institute for Regenerative Medicine.

## Author contributions

T.L.D., E.L.R., A.A.G., and A.T. conceptualized this study and supervised all experiments and data analysis. A.T., N.A., J.L.P.M., K.D., M.P.F., and J.V. performed experiments. A.T., N.A., K.B., N.L., A.M., T.W., and M.P.F. processed and analyzed data. All authors contributed to data interpretation. A.T. wrote the original draft of the manuscript. T.L.D., E.L.R., A.A.G., and A.T. reviewed and edited the manuscript.

## Supplemental Figures

**Figure 1.**
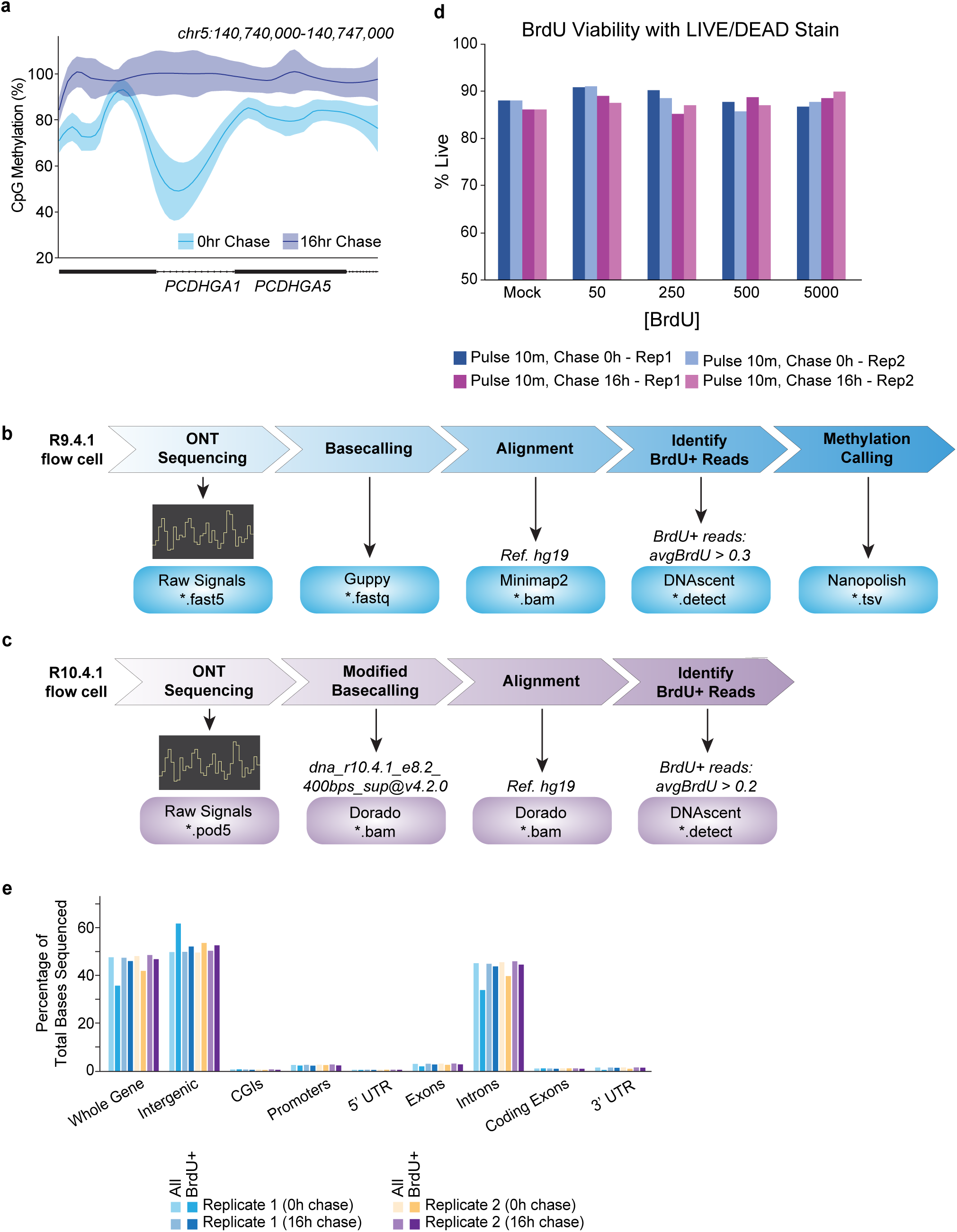
Capturing the temporal dynamics of CpG methylation via long read sequencing. **a)** Representative browser track depicting bulk CpG methylation percentage across BrdU^+^ reads (generated via Repli-BS) that are derived from the 0h chase (nascent) and 16h chase (mature). **b-c)** Bioinformatics analysis pipelines for identifying BrdU-containing reads and calling CpG methylation for **b)** R9.4.1 and **c)** R10.4.1 flow cells from Oxford Nanopore Technologies. **d)** Bar plot depicting % live cells following treatment of HUES64 hESCs with different BrdU concentrations and chase times. Data is shown for 2 biological replicates and was measured through LIVE/DEAD Stain. **e)** Bar plot displaying percentage of total bases sequenced in BrdU-containing reads vs all reads that overlap with annotated genomic regions. Data is shown for both chase timepoints across 2 biological replicates.

**Figure 2.**
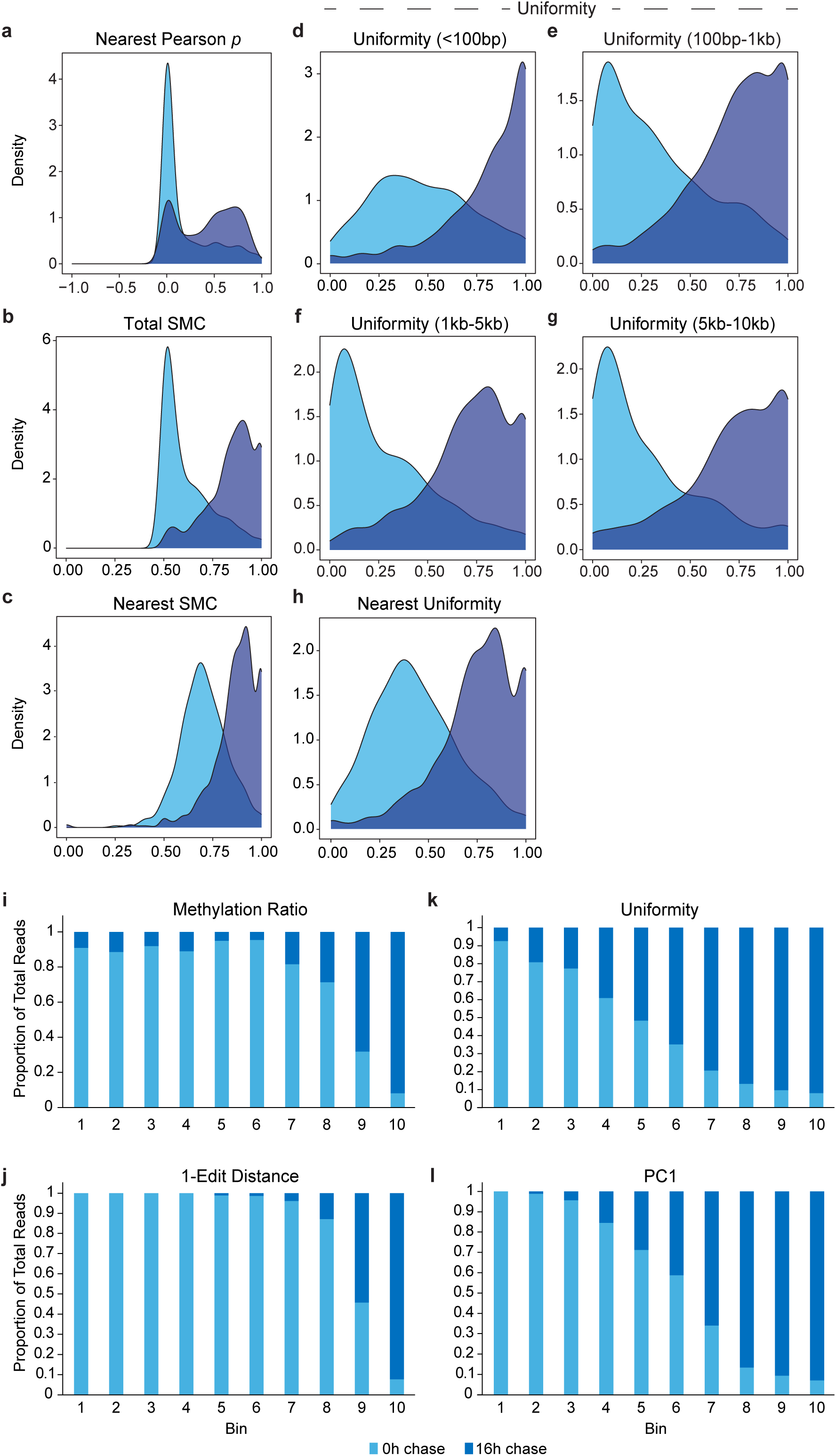
Separation of nascent vs mature chromatin via individual read-level metrics. **a-h)** Distributions of **a)** nearest-neighbor Pearson correlation p values, **b)** simple matching coefficient (SMC) across total reads, **c)** nearest-neighbor SMC, **d)** uniformity of CpGs <100bp apart, **e)** uniformity of CpGs 100bp-1kb apart, **f)** uniformity of CpGs 1kb-5kb apart, **g)** uniformity of CpGs 5kb-10kb apart, and **h)** nearest-neighbor uniformity for reads derived from the 0h (n = 848 reads) and 16h (n = 975 reads) chase timepoints. **i-l)** Stacked bar plots depicting the proportion of BrdU-labeled reads within each **i)** methylation ratio, **j)** 1-edit distance, **k)** uniformity, or **l)** PC1 bin that are derived from the 0h chase vs 16h chase. For each read-level metric, all possible values were ordered least to greatest and divided into 10 bins of equal intervals.

**Figure 3.**
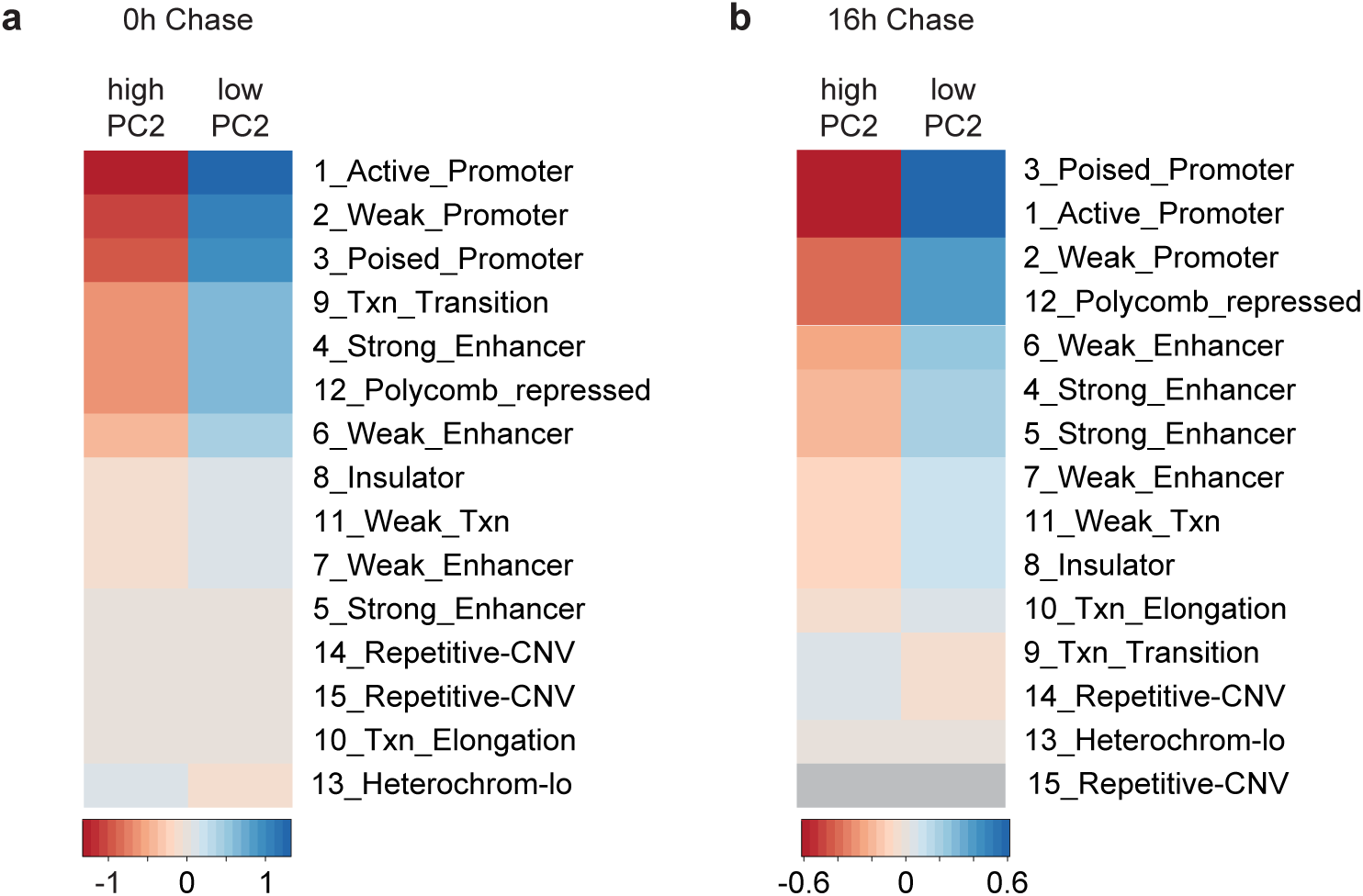
Enrichment analyses of labeled and unlabeled temporally stagnant reads. **a-b)** Heatmap depicting enrichment of high PC2 and low PC2 labeled read populations from the **a)** 0h chase time point and **b)** 16h chase time point for certain genomic regions annotated by chromHMM.

**Figure 4.**
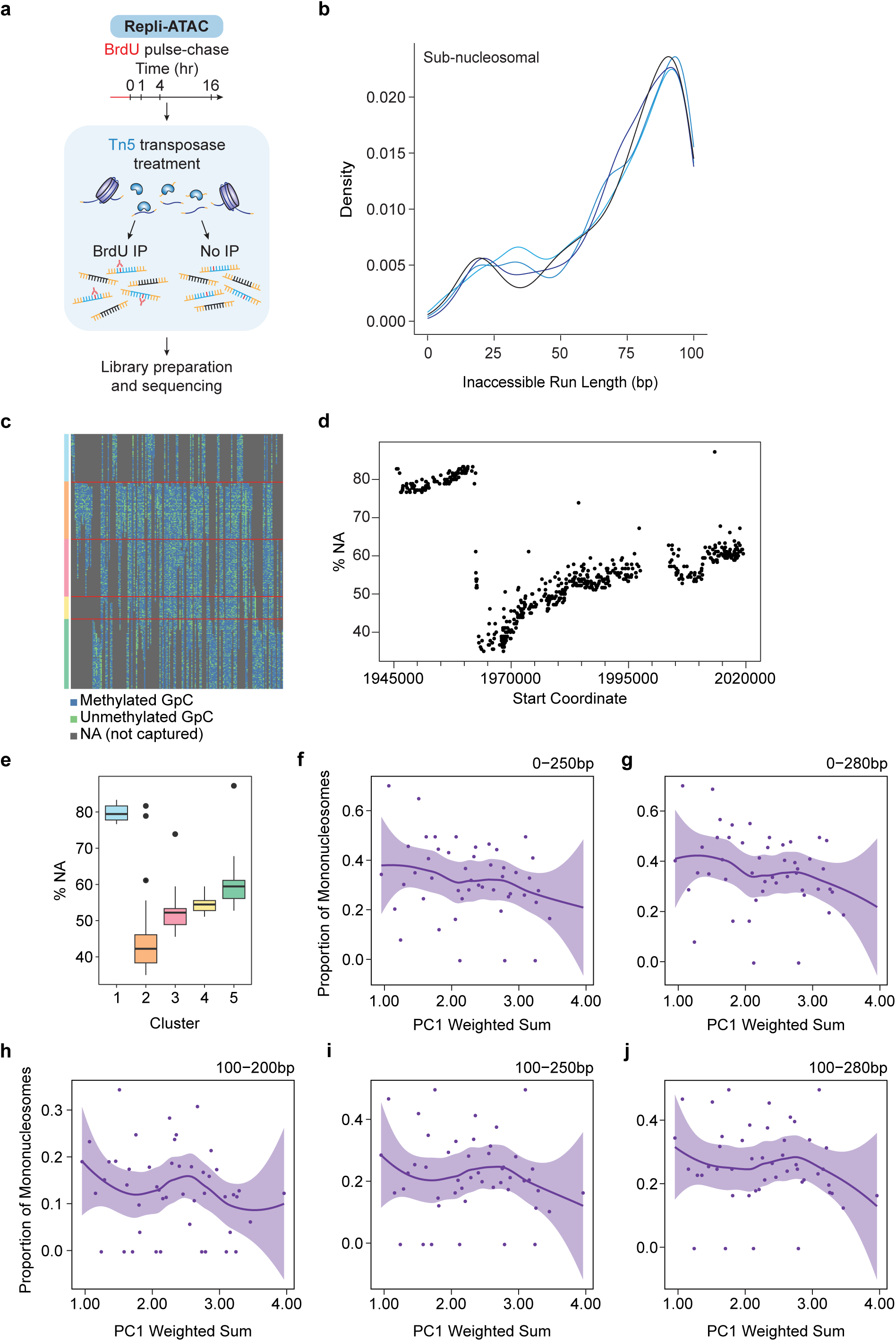
Temporal profiling of nucleosome occupancy using MPATH and publicly available NOMe-seq data. **a)** Schematic illustrating the Repli-ATAC sequencing workflow in HUES64 hESCs. **b)** Density plots showing the distributions of sub-nucleosomal inaccessible run lengths (<100 bp) for 10 kb fragments that fall into each of the PC1 bins (T1-T4). **c)** Heatmap depicting the coverage of GpCs across methylation pseudotime with single-molecule resolution. Only data from the 73kb CRISPR-Cas9 targeted region were used. Only GpCs captured at least 30x are shown (n = 557). Each row represents an individual GpC and each column represents an individual read. Reads are ordered left to right by PC1 weighted sum value (180 reads total) and GpCs are linearly ordered top to bottom. GpCs are colored blue (methylated) or green (unmethylated) if captured by a read and gray if uncaptured by a read. GpCs are clustered based off of their coverage. **d)** Scatter plot depicting the coverage (percentage of reads that are NA for a given GpC) of individual GpCs across the genomic coordinates captured by the 73kb CRISPR-Cas9 targeted region. **e)** Box plot showing the distributions of coverage (percentage of reads that are NA for a given Gaps) for GpCs in Clusters 1 (n = 105), 2 (n = 124), 3 (n = 126), 4 (n = 49), and 5 (n = 153). **f-j)** Scatter plots and Loess smooth curves depicting the proportion of inaccessible run lengths from individual 10 kb fragments that are **f)** 0-250bp long, **g)** 0-280bp long, **h)** 100-200bp long, **i)** 100-250bp long, and **j)** 100-280bp long, plotted across the PC1 weighted sum of the 10 kb fragment. Only 10 kb fragments from Cluster 5 are shown.

