## Supplemental Note 1 for "Methylation pseudotime analysis for label-free profiling of the temporal chromatin landscape with long-read sequencing"

### Supplementary Note 1

### Read-Level Metrics Equations

A single DNA read, and a genome location matched WGBS counterpart, were defined as 1D arrays *x* and *w*, respectively, with equal length *n*:

$$x = [x_1, x_2, \dots\ x_n]$$

$$w = [w_1, w_2, \dots\ w_n]$$

composed of elements representing CpG sites' methylation status within the set:

$$x_i = \{ \quad 0,\ \ unmethylated\ 1,\ \ methylated$$

$$w_i = \{1 \ \geq\ w\ \geq 0, bulk\ methylation\ level\ w \in\ R$$

for site *i*, and number of CpG sites *n* captured in the read:

$$n = \{n\ >\ 0\ n\ \in\ Z$$

The locations of the CpG sites in the genome were defined as a 1D array *r*

$$r_i = [r_1, r_2, \dots\ r_n]$$

within the set:

$$r = \{r\ >\ 0\ r\ \in\ Z$$

#### Methylation Ratio

The mean methylation status of a given read, *x*, was calculated via

$$\underline{x} = \frac{1}{n} \sum_{i=1}^{n} \quad x_i$$

#### Edit Distance

The edit distance for a given read, *x*, was calculated in reference to *w* via

$$D = \frac{1}{n} \sum_{i=1}^{n} \quad (x - w)^2{}_i$$

#### Correlation and Related Metrics

Pearson correlation and associated p values were computed using the standard approach.

Consider two lists of methylation attributes taken from a single read, labeled $V_1$ and $V_2$. The lists may be constructed, for example, as nearest-neighbor pairs. In this case, from the read array *x*:

$$V_1 = [x_1, x_2, \dots\ x_{n-1}]$$

$$V_2 = [x_2, x_3, \dots\ x_n]$$

Then the metrics are defined as follows:

#### Simple Matching Coefficient

For lists $V_1$ and $V_2$, Simple Matching Coefficient refers to the proportion of CpG pairs that are matched in methylation status:

$$SMC = \frac{M_{11} + M_{00}}{M_{11} + M_{10} + M_{01} + M_{00}}$$

where M represents the number of CpG pairs {$V_1$, $V_2$} in which both CpGs are methylated ($M_{11}$), unmethylated ($M_{00}$), or unmatched in their methylation statuses ($M_{10}$ or $M_{01}$). This measure is useful for binary data.

#### Uniformity

A measure related to the SMC, which we refer to as “Uniformity”, is given by:

$$Uniformity = \frac{M_{11} + M_{00}}{M_{11} + M_{10} + M_{01} + M_{00}} - \frac{M_{10} + M_{01}}{M_{11} + M_{10} + M_{01} + M_{00}}$$

That is, it refers to the proportion of pairs that are matched in methylation status minus the proportion of pairs that are not matched in methylation status. Note that this means the Uniformity measure can vary between -1 and 1.

#### Functions of Genomic-Distance or “Total” Reads

All of the above measures can be computed as functions of genomic distance. To this end, pair-wise absolute-value distances, $|r_i - r_j|$, among every CpG on a given read are computed. Pairs are then binned according to distance bins (1, 100, 1000, 5000, 10000) bp. In this way, lists {$V_1$, $V_2$} are constructed as a function of distance, and the above measures may be computed for CpG pairs whose intervening distances falls within one of the aforementioned distance bins.

Additionally, the “Total” metric (for example, Uniformity) may be computed across the entire read, where $V_1$ and $V_2$ contain every pair of CpGs from the whole read.

### Other Equations

#### PC1 Weighted Sum

The weighted sum of PC1 values for a given read is defined as the sum of the proportions of its 10-kb read fragments assigned to each weighted PC1 bin and is otherwise referred to as the PC1 Weighted Sum. Read fragments assigned to T1, T2, T3, T4 are assigned weights of 1, 2, 3, and 4, respectively. The equation is as follows:

$$Weighted\ Sum\ of\ PC1 = 1a + 2b + 3c + 4d$$

where *a*, *b*, *c*, and *d* refer to the proportions of read fragments comprising a given read that were assigned to T1, T2, T3, and T4, respectively.
